# Phage Mu Utilizes the Sliding Clamp for Late Gene Transcription

**DOI:** 10.1101/2025.01.04.631326

**Authors:** Khang Ho, Rasika M. Harshey

**Author notes:** Correspondence to either author.

## Abstract

MuC, along with core RNAP and σ^70^, is required for transcription of late genes of phage Mu. We observed that overexpression of MuC is lethal in *E. coli* and that host replication was over-initiating under these conditions. Suppressors of MuC lethality occurred in dnaA, diaA, and dnaX, genes involved in initiation of DNA replication. We hypothesized that lethality was likely due to increase in ATP-DnaA levels, which are normally kept in check by Hda-DnaN (β clamp). We noticed that MuC harbors a well-defined motif for interaction with the β clamp. We provide experimental support for this interaction and show that disrupting it through mutations in the clamp-binding motif on MuC, expressed from both plasmid and the phage genome, abrogated transcription from a MuC-dependent promoter as well as production of viable phage, respectively. Furthermore, we show that inactivation of β-clamp specifically inactivates C-dependent transcription but not σ^70^-dependent transcription. We conclude that MuC requires the b sliding clamp for transcribing late phage genes. To the best of our knowledge, this is the first demonstration of the involvement of the clamp, a processivity factor essential for DNA replication, for transcription by *E. coli* RNAP.

## Introduction

Phage Mu employs a unique mechanism of replication by transposition [1]. The transposition reaction of the 37kbp Mu genome is carried out by two phage-encoded proteins: MuA, the transposase, and MuB, an AAA+ family protein involved in target capture, as well as two host proteins: IHF and HU. MuA recognizes and binds to two Mu ends, Left and Right, that define limits of the phage [2]. The synapsis of Mu ends is assisted by IHF and HU, which bind and bend DNA to facilitate different transactions on the DNA. MuB non-specifically binds to DNA, forming filaments [3,4]. Target recognition is achieved by the ability of MuA to stimulate the ATPase activity of MuB when they are in proximity, which causes the release MuB from the target DNA [1]. Mu ends and the target DNA are then joined through a nicked-end joining mechanism by MuA [5]. Disassembly of MuA-DNA complex and resolution of the transposition product to generate a new copy of Mu requires a myriad of host factors [6]. Of these host factors, the replication function is carried out by the replicative DNA Polymerase, DNA Pol III.

DNA Pol III is responsible for the replication of the 4.6Mbp genome of *E. coli* [7] in contrast to Pol I, a gap-filling enzyme [8], and translesion (TLS) polymerases for bypassing damaged DNA sites (Pol II, Pol IV, and Pol V) [9]. Pol III consists of 10 distinct proteins, divided into 3 subassemblies: Core Polymerase (α, ε, and θ), Clamp (b), and <u>C</u>lamp <u>L</u>oader <u>C</u>omplex, or CLC, (τ, γ, δ, δ’, χ, and ψ) [10]. DNA synthesis is carried out by the α subunit and ε is the proof-reading subunit (3’-5’ exonuclease activity), while θ serves to stimulate the proof-reading function of ε [10]. The b clamp is the processivity factor that enables Pol III to synthesize long DNA templates (e.g. the *E. coli* genome). In the absence of b, Pol III core polymerase exhibits a processivity of 10bp, approximately that of Pol I [10,11]. Crystal structure of the b clamp reveals the mechanism for processivity: a dimer of the clamp forms a ring-shaped structure that encircles the DNA, thus enabling the core DNA polymerase to stay associated with the DNA during replication [12]. The ring structure of the b clamp is released and loaded onto primer/template junction by the action of the CLC [13]. The minimal pentameric CLC consists of 3 subunits of either τ or γ, one subunit of δ, and one subunit of δ’[14]. In addition to the minimal CLC, two accessory proteins χ and ψ assists in the DNA binding activity of CLC at primer/template junction [15,16]. A cycle of clamp loading and unloading of the CLC is achieved through the binding of ATP and its subsequent hydrolysis [15]. ATP-bound CLC binds and opens the dimeric b clamp. Concurrently, CLC binds to the primer/template junction, which is stabilized by the interaction between χ and single-stranded binding (ssb) protein [15,17]. ATP hydrolysis of the CLC is coupled to the closure of the b clamp ring and release of the CLC from the bound DNA [15].

The clamp is also involved in controlling the rate of replication initiation in *E. coli*. DnaA controls replication initiation at the origin of replication (*oriC*) by binding DnaA boxes at *oriC*, promotes melting of the duplex and subsequent assembly of DnaB, the helicase, the primase, DnaG, and eventually Pol III [18]. Replication initiation by DnaA is regulated by its nucleotide bound state, with only ATP-DnaA being proficient in replication initiation [19]. One mechanism to control the nucleotide bound state of DnaA is known as <u>R</u>egulatory Inhibition of <u>D</u>na<u>A</u> (RIDA). RIDA requires Hda, a homologue of DnaA, as well as the clamp [20–22]. Interaction between the clamp and Hda enables Hda to stimulate the ATP-hydrolysis activity of DnaA, resulting in a decrease in ATP-DnaA levels [23]. RIDA and other mechanisms involved in regenerating ATP-DnaA (stimulating the release or hydrolysis of ATP from DnaA), work in concert to ensure timely replication initiation [18].

Other systems have also employed the clamp for their specific activity. The eukaryotic sliding clamp, PCNA (<u>P</u>roliferating <u>C</u>ell <u>N</u>uclear <u>A</u>ntigen) [24], in addition to its role in DNA replication and repair, is co-opted by LINEs for retro transposition [25]. The bacterial transposon Tn7 interacts with the clamp through its TnsE subunit for target selection [26]. Of particular interest is the coupling of replication to late gene transcription of phage T4 through the T4-encoded sliding clamp [27]. T4 late gene transcription requires host core RNA Polymerase (RNAP), two RNAP binding proteins gp33 and gp55 [27–29], which together could be thought of as a split sigma factor, the T4 clamp (gp45), which participates in both T4 DNA replication as well as activation of late gene transcription [30], and the T4 clamp loader complex (gp44/gp62) [27], though this requirement could be bypassed *in vitro* with a crowding agent such as PEG [31]. DNA structures transiently generated by T4 replication in the form of nicks or primer/template junctions promote loading of the clamp by the gp44/gp62 clamp loader [27]. Gp55/Gp33 bound RNAP is activated upon contact between Gp55 acidic C-terminal and Gp45 [32], thus coupling replication and late gene activation in T4 phage.

The phage Mu transcription factor MuC is responsible for transcription of Mu late genes [33]. MuC is a 16.5kD polypeptide that binds 4 specific DNA sequences on the Mu genome to activate transcription [34–36]. These sites reveal a bipartite consensus sequence 5’-TTAT-[N_5-6_]-ATAA-3’ [37]. MuC-dependent promoters possess a recognizable -10 sequence but lack a recognizable -35 sequence [34,35]. MuC requires σ^70^ along with core RNAP (a_2_, b, b’, w) to activate transcription [36,38]. However, it does not require the C-terminal domain of the α subunit nor the C-terminal domain of σ^70^ [38].

In this report, we show that MuC interacts with the clamp, and that this interaction is required for its transcriptional activity. This discovery came about serendipitously when we observed that overexpression of MuC is toxic. Flow cytometry revealed that *E. coli* replication over-initiates under these conditions, indicative of ATP-DnaA accumulation. Suppressors of this phenotype occurred in *dnaA*, *diaA*, and *dnaX*. These and other observations suggested that overexpression of MuC was disrupting RIDA, either through the clamp or through Hda. We found that MuC harbors a well-defined clamp-binding motif required for MuC-dependent transcription, and that inactivating the clamp specifically abolishes transcription from a MuC-dependent promoter, but not from any σ^70^ promoter. We conclude that MuC is a clamp-dependent transcription factor.

## Results

### Overexpression of MuC is toxic and is suppressed by genes involved in replication

MuC, which is required for transcription of four phage late-gene promoters (p*lys*, p*P*, p*I*, and p*mom* [35]) has an unusual Ribosome Binding Site (RBS) missing the 6-7 bp spacer between the Shine-Dalgarno sequence and the start codon (Fig. 1A). Expression of MuC both *in vitro* and *in vivo* was reported to be inefficient [34,35]. We wondered whether the atypical RBS was a regulatory sequence, requiring a phage product for efficient expression. To test this hypothesis, we cloned MuC under the inducible (by anhydrotetracycline or aTc), Tet promoter and compared MuC expression by aTc induction to its expression when the Mu prophage was induced (by heat-inactivation of its temperature-sensitive repressor). However, we were surprised to find that induction of MuC by aTc was toxic (Fig. 1B), resulting in a 3-log loss of CFU (colony forming units). MuC overexpression has been reported previously without this phenotype [35,39]. We decided to investigate the mechanism of this toxicity by looking for suppressors. To avoid isolation of mutants that simply mutated MuC, we ascertained MuC functionality through its ability to activate transcription of the *mom* promoter (p*mom*) fused to *lacZ* on a compatible plasmid, and selected for blue survivors of aTc induction on plates supplemented with X-gal. We isolated 14 mutants and performed whole genome sequencing to identify mutations in these strains. The data are presented in Table 1 (see Methods for cutoff of hits shown).

**Figure 1.**
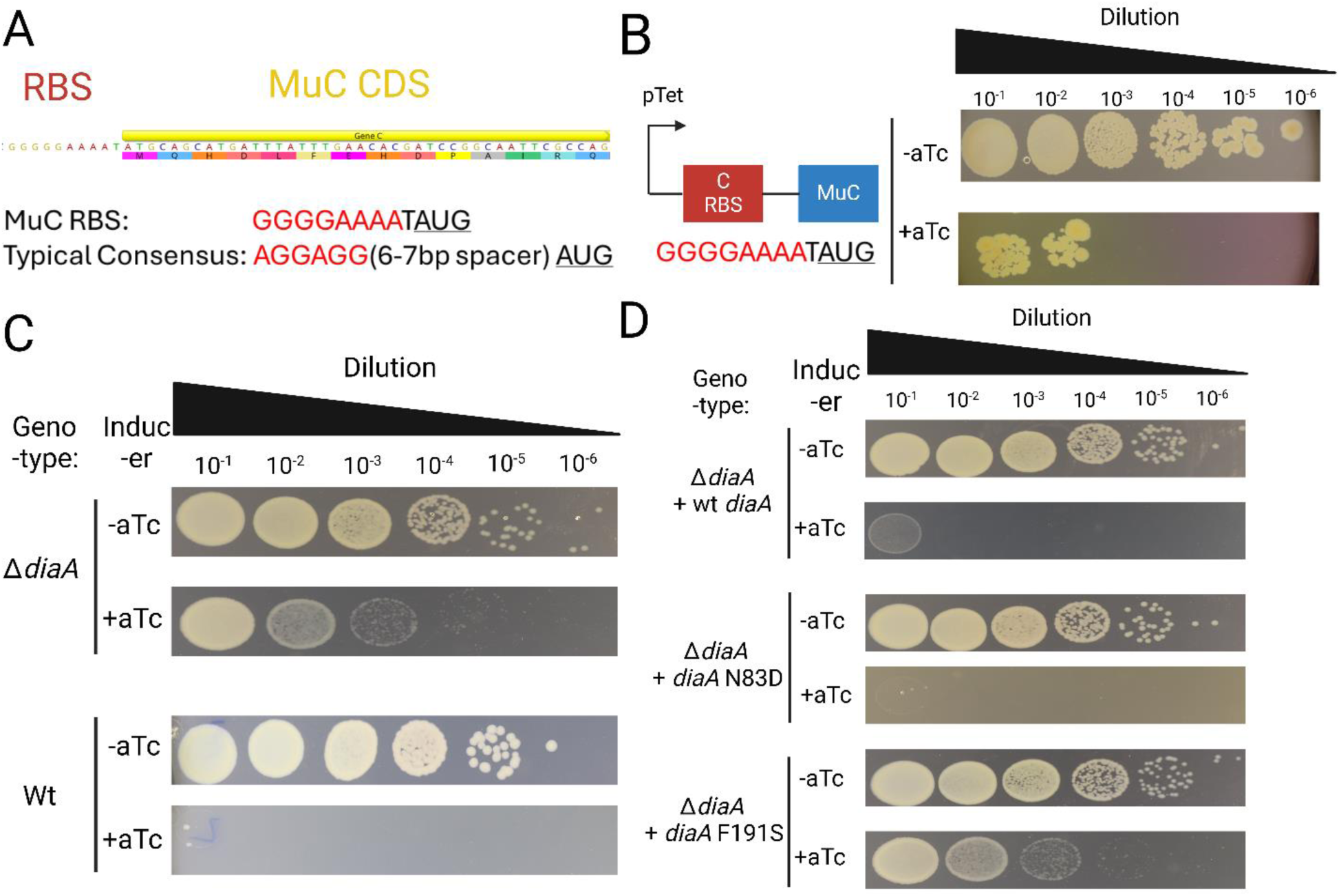
Toxicity of MuC overexpression and its suppression by DiaA. A) The ribosome binding site (RBS) of MuC. The coding sequence (CDS) is yellow. Single letter amino acids are shown below the coding sequence. The RBS is in red. B) MuC expression from a pTet promoter is toxic when induced by aTc. C) Partial rescue of MuC overexpression toxicity (see B) in Δ*diaA* background. D) Effect of known *diaA* variants on MuC toxicity (see text).

**Table 1.** Suppressors of MuC Toxicity and their associated mutations. Codon number indicates the position of the resulting change in the amino acid sequence. Variant frequency is the frequency of reads with indicated allele. See Material and methods for cut-off thresholds.

| Strain ID | Amino Acid Change | Gene Name | Codon Number | Variant Frequency |
| --- | --- | --- | --- | --- |
| 1 | Q -> K | *flhB* | 25 | 100.00% |
|  | G -> A | *diaA* | 148 | 100.00% |
| 2 | Q -> K | *flhB* | 25 | 99.40% |
|  | G -> A | *diaA* | 148 | 100.00% |
| 3 | Frame shift | *sapB* | 228 | 100.00% |
| 4 | L -> * | *diaA* | 94 | 98.40% |
| 5 | P -> A | *dnaX* | 380 | 100.00% |
|  | L -> * | *diaA* | 94 | 100.00% |
| 6 | A -> P | *zntA* | 675 | 99.50% |
|  | A -> P | *ydhK* | 392 | 100.00% |
|  | L -> * | *diaA* | 94 | 100.00% |
| 7 | A -> P | *yqgE* | 13 | 100.00% |
|  | L -> * | *diaA* | 94 | 99.50% |
| 8 | $\Delta 96-118$ | *dnaA* | N/A | 100.00% |
| 9 | R -> H | *dnaA* | 342 | 100.00% |
| 10 | G -> A | *fiu* | 217 | 99.60% |
| | $\Delta 96-118$ | *dnaA* | 96 | 100.00% |
| 13 | Frame shift | *sapF* | 50 | 100.00% |
| 14 | L -> P | *trkA* | 6 | 98.60% |
| | $\Delta 419-425$ | *rsmB* | 425 | 99.50% |

We were surprised to find that the suppressors mapped to genes involved in DNA replication. Two distinct mutations were recovered in *dnaA*: a small deletion of the linker domain II (Δ96-118), and a missense mutation in the ATP-binding domain III (R342H) [40]. We will refer to the domain II and III mutants as *dnaACr8* (<u>C</u>-resistant) and *dnaACr9*, respectively. Encouragingly, we observed two mutations in *diaA*, a known partner that promotes DnaA assembly on to *oriC* [41,42]: a premature stop codon at position 94, predicted to result in a truncation that produces one-half the protein, and a missense mutation (G148A). We also observed a missense mutation in *dnaX* (P380A), which affects the τ/γ subunit of the CLC.

Other mutations recovered included the Sap exporter, involved in putrescine export [43], a gene of unknown function (*yqjK*), a gene involved in DNA damage response (*yqgE*) [44]. Of note is a pair of mutations that affect two adjacent genes: *trkA*, a potassium importer [45], and *rsmB*, an RNA methyltransferase [46]. We sought to determine which of these mutant alleles could provide resistance against MuC toxicity by providing each on a separate plasmid and assaying for rescue of MuC toxicity. However, neither did, so we will not discuss them further.

We next sought to verify that the mutations in DNA replication functions we found were related to MuC toxicity. DnaA is required for replication initiation, and DnaX is a component of Pol III, rendering their inactivation difficult, so we chose *diaA* for deletion, since it has been shown to be non-essential [41]. Deletion of *diaA* permitted the growth of cells when MuC expression was induced (Fig. 1C, top panel) as compared to that of the Wt strain (Fig. 1C, bottom panel). Conversely, we were able to complement the resistance to MuC toxicity by providing *diaA* in *trans* on a plasmid, restoring sensitivity to MuC toxicity (Fig. 1D, top panel). DiaA has been shown to oligomerize and interact with DnaA to promote loading specific loading of ATP-DnaA at *oriC* [42]. Point mutations in *diaA* that perturb DiaA oligomerization or its interaction with DnaA and the consequences thereof upon replication initiation have been described [42]. To better understand the mechanism of *ΔdiaA* resistance to MuC toxicity, we introduced these point mutations into our *diaA* expression vector in a Δ*diaA* background and found that the mutation that interferes with DiaA oligomerization (N83D) restored MuC sensitivity to a Δ*diaA* background, much like Wt DiaA (Fig. 1D, middle panel). By contrast, *diaA* F191S, which is defective for DnaA binding showed resistance to MuC like the Δ*diaA* strain (compare Fig. 1D, bottom panel to Fig. 1C top panel). We interpret these results to mean that in the absence of DiaA, it is the lack of interaction between DiaA and DnaA that allows for MuC resistance. Therefore, we hypothesize that the defect caused by MuC overexpression affects the function of DnaA, either directly or indirectly.

Conversely, might DnaA affect the function of MuC, and, by extension, inhibit late gene transcription of Mu? To address this, we placed Wt DnaA or DnaA alelles isolated from the suppressor screen (Table 1) under an inducible promoter and monitored: (1) lysis curve of a Mu lysogen and (2) transcription from p*mom*, the MuC-dependent promoter. The lysis curves showed delayed lysis upon induction of Wt DnaA but not of *dnaACr8*, with *dnaACr9* exhibiting an in-between phenotype (Fig. S1 A,C,E). Expectedly, p*mom* transcription was decreased with Wt DnaA (Fig. S1B), with the two *dna* alleles behaving differently: *dnaACr8* actually stimulating p*mom* (Fig. S1D) by about 2-fold, and *dnaACr9* having no effect (Fig. S1F). This result was unexpected since DnaA was shown to be not required for Mu development [47]. We have not pursued here how DnaA might affect Mu transcription.

In summary, the results in this section suggest that MuC toxicity is a consequence of perturbation in the replication machinery, likely through DnaA. The effect of DnaA variants identified through the suppressor screen on MuC transcription also suggests that MuC might need some component of the replication apparatus for its transcription.

### MuC Overexpression Causes Replication Over-initiation through Accumulation of ATP-DnaA

We next sought to elucidate the effect of MuC overexpression on *E. coli* replication initiation. To do so, we performed flow-cytometry and labelled genomic DNA with Sytox green. Most cells grown in LB without overexpression of MuC contain 4 chromosomes, and a small population contains 8 chromosomes (Fig. 2A, green curve). With MuC overexpression, we observed a loss of discrete chromosome copy number, accompanied by an increase in DNA levels in cells (Fig. 2A, right-shifted blue curve) relative to uninduced control. This is indicative of over-initiation of replication: replication re-initiates before Pol III can finish the ongoing round, resulting in a loss of discrete chromosome numbers while increasing the amount of DNA content in cells. We then examined the DNA content of Δ*diaA* with or without MuC overexpression to determine if absence of *diaA* would suppresses the observed increase in DNA content found with MuC in Wt. In the absence of MuC induction, Δ*diaA* cells have a variable number of chromosome copies (Figure 2A, red curve), ranging from 1 to 4, consistent with previous findings [41]. With MuC induction, Δ*diaA* cells still show an increase in DNA content (Fig. 2A, orange curve) and loss of discrete chromsome copies, but not as severe as that of Wt (i.e Δ*diaA* curve is left-shifted compared to blue MuC curve, indicating less DNA content). We conclude that MuC overexpression causes over-initiation of replication.

**Figure 2.**
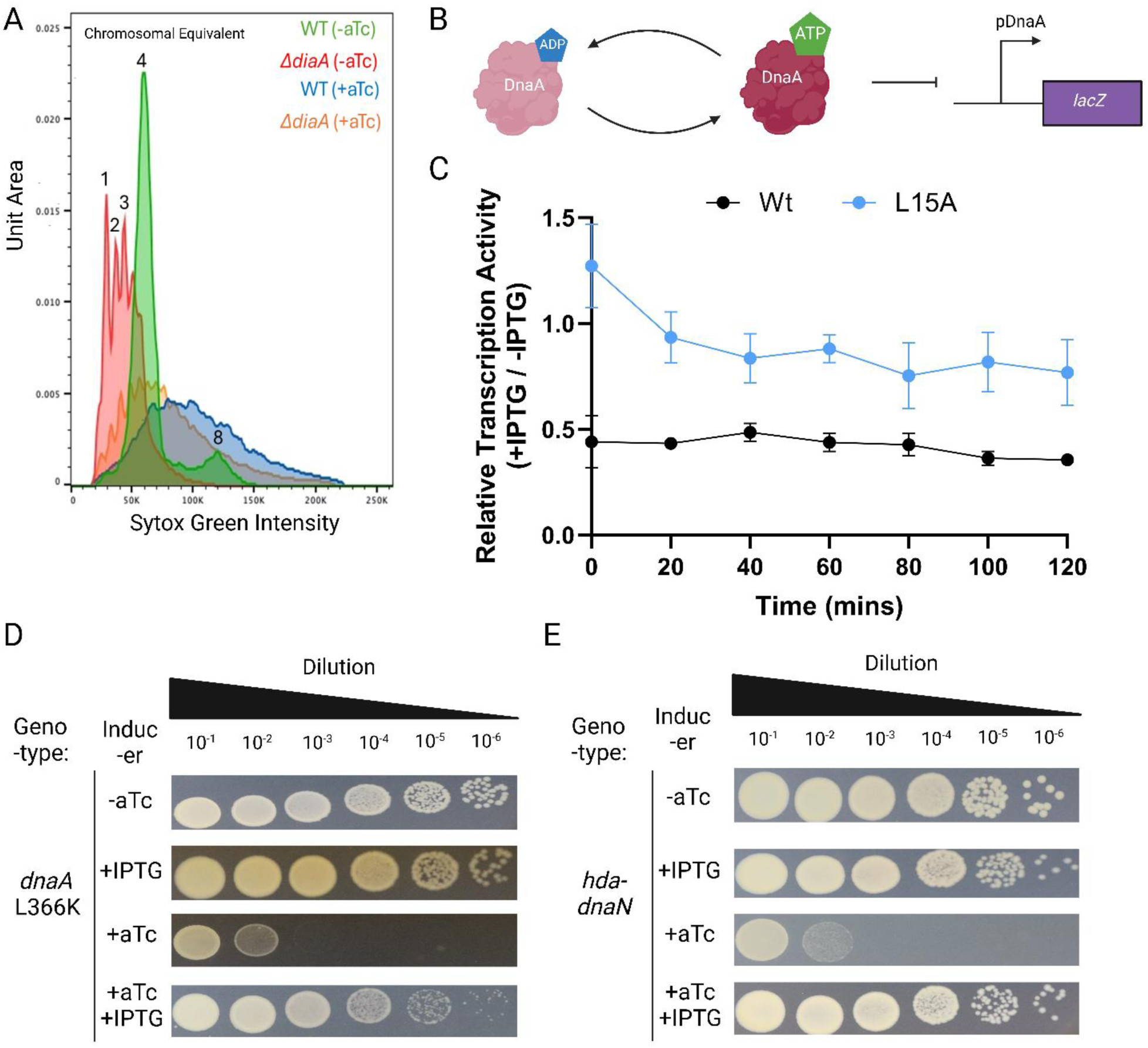
Replication over-initiation caused by MuC overexpression and its suppression. A) Flow cytometry MuC overexpressing strain. DNA content, indicated by with Sytox green intensity, and chromosomal equivalents (number above indicated peaks) are shown. The chromosomal equivalent is determined by performing the same assay on stationary phase cell (overnight culture). B) Scheme to determine ATP-DnaA level through its repression of p*dnaA*. C) Effect of MuC expression on repression of p*dnaA*. The error shown is from 3 biological replicates. D) Rescue of MuC toxicity by expression of DnaA L366K. E) Rescue of MuC toxicity by co-expression of *hda* and *dnaN* clamp).

Replication initiation in *E. coli* is controlled by the nucleotide-bound state of DNA [18,19]. DnaA-ATP forms spiral-like oligomers and initiate replication by melting *oriC* and promoting stepwise assembly of the Pol III [48], but DnaA-ADP is not proficient in these activities. Strains compromised for RIDA (Δ*hda*) have been shown to accumulate ATP-DnaA [21]. We hypothesized that MuC overexpression causes over initiation by increasing the amount of ATP-DnaA. To test this, we exploited the ability of DnaA to autoregulate its transcription: DnaA-ATP binds and represses transcription from its promoter, while DnaA-ADP is capable of binding but not repression [49]. We fused the promoter region of DnaA that is subjected to ATP-DnaA repression to *lacZ*, enabling us to determine ATP-DnaA level through b-galactosidase activity. We expected that high levels of ATP-DnaA would repress transcription from p*dnaA*, resulting in low b-galactosidase activity, and vice versa (Fig. 2B). With this reporter system, we observed that with MuC overexpression (Wt), b-galactosidase activity decreased compared to the uninduced control by 58% on average across time points (Fig. 2C, black line represents ratio of induced to uninduced), consistent with the hypothesis that MuC overexpression results in accumulation of ATP-DnaA. Next, to verify that the increased ATP-DnaA levels, approximated by decrease in *lacZ* reporter, is linked to MuC toxicity, we used a MuC L15A mutant that abolished MuC toxicity without abolishing its transcriptional activity (Fig. S2A-C). We observed that overexpression of MuC L15A produces a modest repression of *lacZ* of 90% of uninduced activity on average across time points (Fig. 2C, blue line), supporting our hypothesis that MuC toxicity is linked to accumulation of ATP-DnaA levels.

To support our hypothesis that ATP-DnaA is elevated upon MuC overexpression we provided other conditions known to regulate DnaA-ATP levels, with the expectation that they would rescue the MuC toxicity. For example, acid phospholipid is known to facilitate nucleotide exchange of DnaA *in vitro* [50,51]. *In vivo*, DnaA mutations in an amphipathic helix suppress a block in the first committed step of phospholipid biosynthesis through *pgsA* [52]. One mutant, DnaA L366K has been shown to have altered interaction with phospholipid without any significant change in its affinity for ATP or *oriC* binding *in vitro* [53,54]. We hypothesized that DnaA L366K, which could bypass the requirement for lipid in nucleotide exchange, suggesting that it is a variant that promotes release of ATP from DnaA, could rescue the accumulation of ATP-dnaA caused by MuC overexpression. We co-expressed either Wt DnaA or DnaA L366K, inducible with IPTG, with aTc inducible MuC, and assessed the ability of DnaA variants to rescue MuC toxicity. With induction of DnaA L366K, we observed rescue of MuC toxicity (Figure 2D, +aTc and +IPTG panel), but not with Wt DnaA (Fig. S2D, +aTc and +IPTG panel). Without IPTG induction, DnaA L366K does not alleviate the effect of MuC toxicity (Figure 2D, +aTc and -IPTG panel). Since phospholipid-dependent nucleotide exchange is not specific to ATP-DnaA, but rather both the ATP and ADP form, we examined whether the clamp-Hda complex mediates RIDA by hydrolyzing ATP-DnaA to its ADP form. Hda, <u>H</u>omologue of <u>D</u>na<u>A</u>, was identified a suppressor of the hyper-initation phenotype of DnaAcos (cold sensitive) [21]. Hda is homologous to domain III, the ATP-binding domain, of DnaA, and promotes hydrolysis of ATP-DnaA in conjunction with the clamp [55]. We introduced either *hda, dnaN* (clamp) individually or together and assessed their ability to rescue MuC toxicity. Expression of either *hda* or *dnaN* alone did not rescue MuC toxicity (Fig. S2E, +aTc +IPTG panel). However, co-expression of *hda* and *dnaN* provided full rescue of MuC toxicity (Fig. 2E, +aTc and +IPTG panel). This suggests that the ATP-hydrolysis activity of the Hda-DnaN complex could rescue MuC toxicity, consistent with our hypothesis that MuC overexpression results in ATP-DnaA accumulation.

Since flowcytometry (Fig. 2A) was performed on cells grown in liquid culture, we sought to verify that the observed rescue of MuC toxicity with either DnaA L366K or Hda/Clamp expression also occurs in liquid growth. We first verified that induction of MuC results in slow growth in liquid, though not complete inhibition (Fig. S2F). We also observed that the Δ*dia* strain permitted partial rescue, consistent with data from plate assay (Fig. S2F). We recapitulated DnaA L366K rescue of MuC toxicity in liquid growth (Fig. S2G). In liquid, even leaky expression of DnaA L366K is sufficient to rescue growth (Fig. S2G), and we observed full rescue with addition of IPTG (Fig. S2E). Curiously, we were unable to rescue MuC toxicity by *hda* and *dnaN* co-expression in liquid (Fig. S2H).

Taken together, we conclude that MuC overexpression causes an accumulation of ATP-DnaA, resulting in replication over-initiation. We reasoned that MuC might be interfering with mechanism(s) that regulate ATP-DnaA levels, rather than interacting with DnaA, due the inability of Wt DnaA overexpression to rescue MuC toxicity. Since we observed rescue of MuC toxicity by providing RIDA, a simple hypothesis is that MuC is interfering with RIDA.

### MuC-Clamp Interaction is Required for Toxicity

Many *E. coli* proteins involved in replcation are known to interact with the clamp, giving rise to a consensus binding motif of 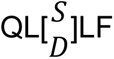 in bacteria [56]. We found that MuC contains a near-consensus clamp binding motif (CBM) at its N-terminus (Fig 3A, top), deviating from the consensus sequence only at the second position (Q<u>H</u>DLF instead of Q<u>L</u>DLF). To determine if MuC interacts with the clamp *in vivo*, we used the bacterial two-hybrid system where interaction between two proteins fused to distinct fragment of *B. pertussis* adenylate cyclase results in production of cyclic-AMP, in turn, stimulating b-galactosidase activity in a *cyaA^-^* strain, which can be detected with X-gal. Fusion of MuC-T25 and T18-DnaN produced positive b-galactosidase activity, indicative of interaction (Fig. 3A, bottom). We verifed that this interaction is dependent on the CBM by constructing point mutants in the CBM and producing quantitative b-galactosidase measurements of the MuC-clamp interaction. Alanine substitution at position 1 (Q2A; numbering of position is according to Fig. 3A, top), produced no significant change in MuC-clamp interaction (Fig. 3A, bottom). L5A mutation abolished MuC-clamp interaction. L15A mutation, identified previously as a less toxic variant of MuC, had a slightly reduced interaction of about 60% of Wt (Fig. 3A, bottom). We also generated a mutation at the non-consensus position of the CBM to produce the consensus sequence (i.e. H3L), with the expectation that this variant would show increased interaction with the clamp. Indeed, the H3L variant shows a 1.6-fold increase in MuC-clamp interaction (Fig. 3A, bottom). We also combined mutations to determine if these mutations elicit synergistic effects on MuC-clamp interactions. Combination of L5A and L15A did not reduce the MuC-clamp interactions more than L5A alone while L5A Q2A combination showed a large decrease (60%) compared to L5A alone (Fig. 3A, bottom). To assess whether any of these mutations affect protein stability, we performed Western Blot on all these variants using a C-terminal FLAG-tagged version. We observed that MuC-FLAG exhibits a slight decrease in MuC toxicity (Fig. S2A), which we judged was not sufficient to affect our interpretation of the tagged MuC stability. We confirmed that most of the MuC variants were produced in similar amounts (Fig. S3A). We note that there was a slight decrease in protein amounts in the L5A and Q2A mutations (15% and 25% compared to Wt, respectively), and a corresponding increase in the H3L variant (25% of Wt). The Q2A L5A variant showed a significant decrease of 57% in protein levels compared to Wt (Fig. S3A).

**Figure 3.**
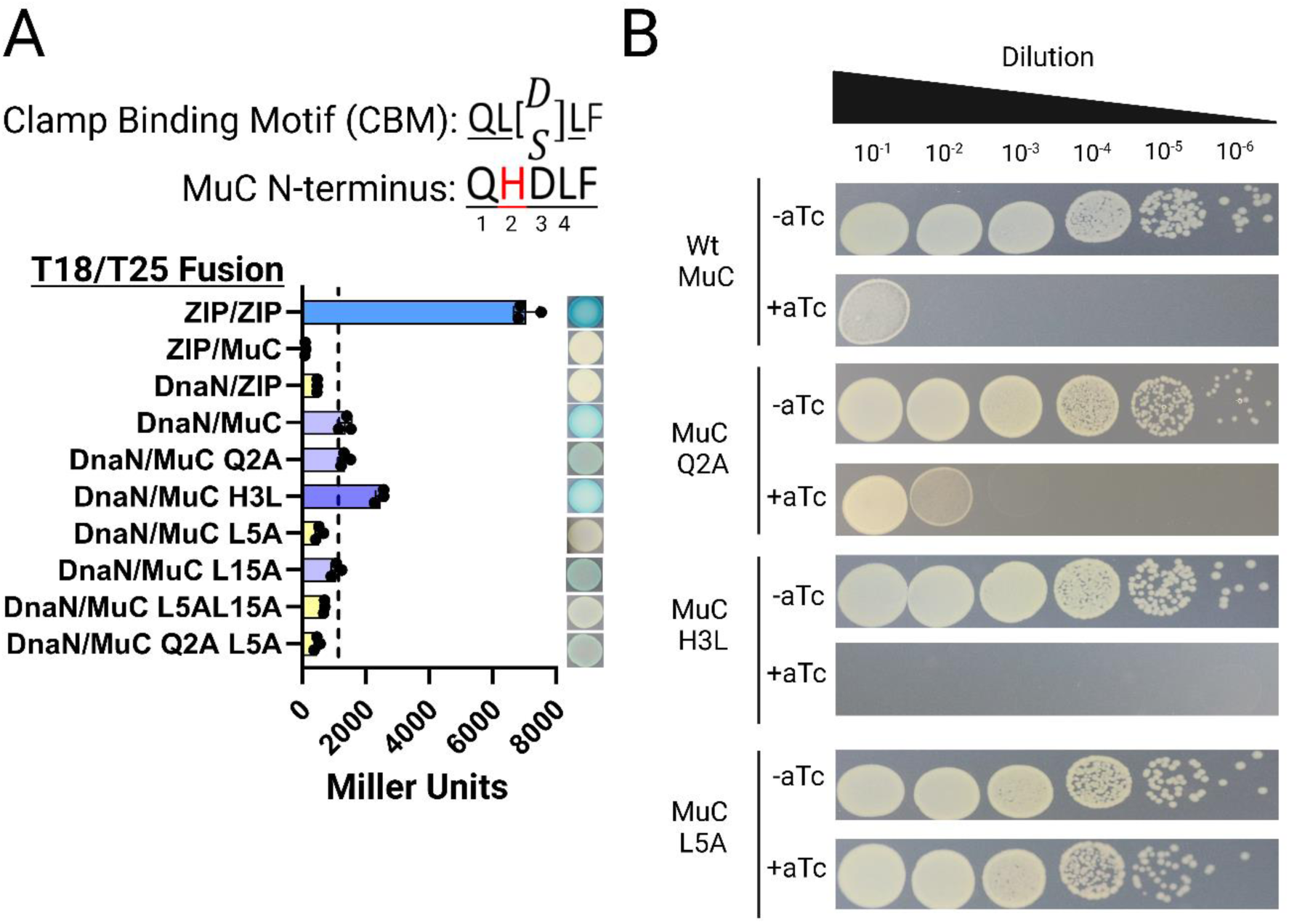
The interaction of MuC with the Clamp is required for its toxicity. A) Top. Motif for clamp binding (CBM) and its present in MuC. Bottom. Measurement of MuC, and its variants, interaction with the Clamp bacterial two hybrid system. Individual data are denoted by black circles. Bar shows the mean value. B) Effect of MuC variants in the CBM on MuC toxicity.

To determine if MuC-clamp interaction is responsible for MuC toxicity, we examined the ability of the CBM mutants to inhibit growth. Q2A mutant was not compromised for toxicity compared to Wt (Fig. 3B). The L5A mutation was completely unable to inhibit growth (Fig. 3B), consistent with this mutant’s inability to interact with the clamp. By contrast, the H3L variant was more effective at growth inhibition than Wt (Fig. 3B), consistent with its increased clamp interaction. The effect of these variants on solid media was recapitulated in liquid (Fig. S3B). Since the clamp is associated with DNA replication, we wondered if the SOS response might be induced during MuC overexpression. However, we did not observe any increase in *sulA* promoter activity (Fig. S3C) or an increase in cell length (Fig. S3D), both indicative of the SOS response.

The transcription factor Mor, that activates the middle operon of Mu that includes MuC [57,58], also possesses a CBM, and has a high sequence similarity to MuC [57]. Mor does have a CBM, albeit smaller than that of MuC, with only the sequence DLF of the consensus 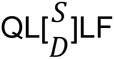 (Fig. S3F, top). We confirmed that Mor overexpression is toxic, albeit to a lesser extent than that of MuC (Figure S3E, bottom), consistent with its smaller CBM. Since Mor/C defines a superfamily of transcription factors [59], we examined 100 other members of this family to determine if they possess a CBM. We found the consensus sequence at of N-terminus of the Mor/C superfamily is Qx[D/E][L/M]F (Fig. S3F), resembling the CBM, the second position being tolerant of substitution reported previously [56]. The high conservation at position 2, 5 and to a lesser extent position 4 and 3 agrees with previous report [56]. We infer from this consensus that clamp binding is a general property of the Mor/MuC superfamily of transcription factors.

We interpret these results to mean that MuC toxicity is a result of its interaction with the clamp. When MuC is overexpressed, it sequesters the clamp through the CBM, preventing the clamp from participating in RIDA.

### MuC Utilizes the Clamp for Transcription

Given that MuC is a transcription factor and that a phage-encoded clamp is known to activate late gene transcription in T4 phage [27], we hypothesized that MuC is utilizing the clamp for stimulating late gene transcription. To test this, we first measured transcription from p*mom* using MuC variants we constructed that show a range of clamp interaction. Like the ability of MuC L5A to inhibit cell growth, the L5A variant is completely defective for p*mom* transcription (Fig. 4A, diamond) compared to Wt (Fig. 4A, circle). The Q2A variant shows a decrease in p*mom* transcription (Fig. 4A, square), ranging from 50% to 60% of the Wt promoter across the time course. By contrast, the H3L variant shows an increase, at a maximum of 130% of the Wt at the final time point (Fig. 4A, triangle). These results are consistent with the degree of interaction of each of these variants with the clamp i.e. MuC transcriptional activity is positively correlated with clamp interaction. Next, to further show that the clamp is essential to MuC transcriptional activity, we assessed the effect of inactivating the clamp on MuC transcription. To do this, we exploited a temperature sensitive allele of *dnaN* (Fig. 4B). We observed that upon shifting to the non-permissive temperature (inactivation of the clamp), MuC transcription decreased with time, reaching 80% reduction in transcriptional activity after 2 hours (Fig. 4C, empty circle) compared to the permissive condition. This decrease did not occur with p*tetA*, σ^70^-depedent promoter (MuC also requires σ^70^ [36]), but, instead, p*tetA* activity increased by 2-fold at 80 minutes and then levelled off (Fig. 4C, empty square). We could complement the decreased MuC transcription due to clamp inactivation by providing Wt *dnaN* in *trans*, which shows a slight stimulation of transcription at non-permissive temperature of a about 1.3-fold (Fig. 4C, filled circle). We did not observe a general stimulation of transcription of p*tetA* promoter by providing Wt *dnaN* in *trans* (Fig. 4C, filled square). Since the clamp is known to require the CLC, we also determined whether the clamp loader is required for MuC transcription using a temperature-sensitive allele of the τ/γ subunit of the CLC. Similar to the clamp, we observed a decrease in MuC transcription upon temperature shift but not that from p*tetA* (Fig. S4A). These results suggest that MuC utilizes the clamp for its transcriptional activity.

**Figure 4.**
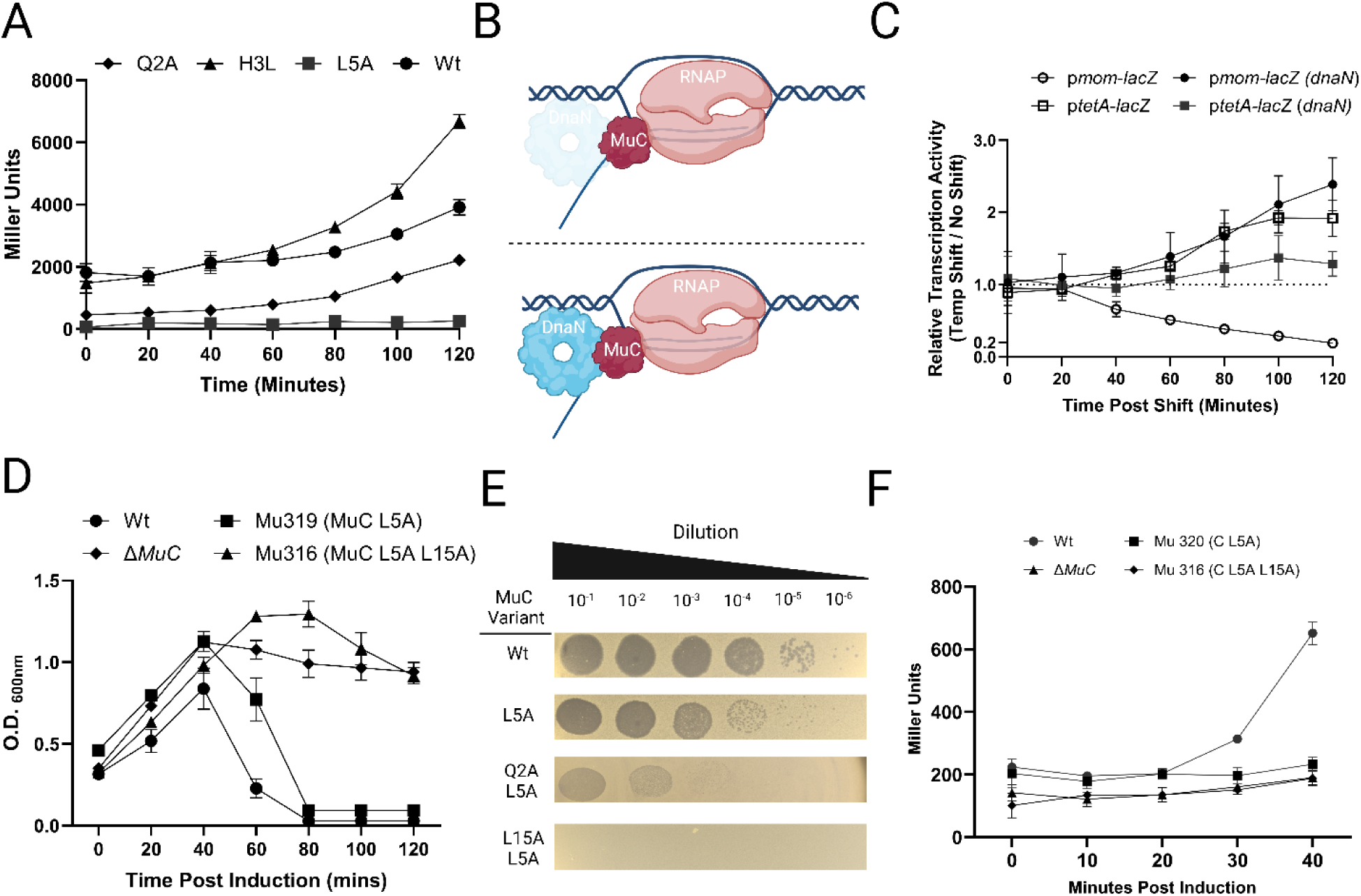
Mu utilizes the clamp for late gene transcription. A) Transcriptional activity of MuC variants. B) Scheme to determine the effect of Clamp inactivation on MuC transcription. C) MuC transcription requires the clamp. D) Lysis curve of MuC variants defective for Clamp interaction. E) Plaquing of Mu phage carrying MuC variants defective for clamp interaction. F) MuC transcription from Mu phage carrying MuC variants defective for clamp interaction.

We next sought to determine if the effect of the clamp we were observing would extend to the Mu phage. We introduced MuC variants CBM mutation(s) into Mu prophages and assayed the ability of these mutant prophages to either plaque or lyse the host cell. Of the single point mutants, only L5A exhibited delay in lysis (Fig. 4D, square) by 20 minutes, while the remaining point mutants showed identical lysis to the Wt (Fig. S4B). Double mutants in the CBM result in more severe delay in lysis: the Q2A L5A showed an 80-minute delay in lysis (Fig. S4B), even though it showed severely lowered stability (Fig. S3A), and the L5A L15A never lysed (Fig. 4D, triangle). This was interesting to us since the Q2A L5A mutant showed severe reduction in MuC levels, while the L5A L15A mutant produced MuC at levels comparable to Wt, yet the L5A Q15A never lysed the host, suggesting that the L5A L15A mutant may be defective for another function or a step downstream of clamp interaction. We also assayed lysate of the mutant prophages for the ability to plaque (chloroform-treated supernatant of the L5A L15A mutant was used, since no lysis was observed). Like the lysis phenotype of the single mutants, only L5A showed a decrease in plaque size (Fig. 4E). The other single point mutants showed no defect in plaquing (Fig. S4C). The double mutant Q2A L5A showed a 3-log decrease in PFU, and the L5A L15A showed no plaquing. We further examined whether the defect we observed in the L5A, Q2A L5A, and L5A L15A mutants are related to late gene transcription by measuring transcription from p*mom* in these lysogens. Indeed, we observed that late gene transcription was defective in all of prophages that contains a L5A mutation, comparable to that of a Δ*MuC* prophage (Fig. 4F). The other mutants showed varied levels of p*mom* transcription (Fig. S4D), consistent with previous MuC result from plasmid expression (Fig. 4A). Since the clamp is often associated with Pol III activity and since Mu requires Pol III for resolution of its transposition products, we initially hypothesized that decreased late gene transcription of the Mu mutants we made is a direct consequence of the Mu replication defect. However, when we measured replication of the L5A L15A mutant, which is completely defective for lysis and transcription, we found that this mutant was as proficient as the Wt for replication (Fig. S4E). We conclude from all these results that Mu utilizes the clamp for its late gene transcription and not for its replication.

## Discussion

We have shown in this study that the phage Mu transcription factor MuC, utilizes the clamp for its transcriptional activity. We first found that MuC overexpression is toxic, and this toxicity is alleviated by mutations in genes involved in replication. We then showed that MuC overexpression causes over-initiation of replication caused by an accumulation of ATP-DnaA. We could suppress MuC toxicity by either overexpressing a DnaA mutant that bypasses the need for lipid in nucleotide-exchange, or by providing components of Hda and the clamp, which constitute the components of RIDA. We demonstrated that MuC toxicity requires the clamp-binding motif on MuC, suggesting that MuC toxicity is a result of disruption of RIDA through the clamp (Fig. 5). We also showed that the MuC-clamp interaction is crucial for Mu late gene transcription.

**Figure 5.**
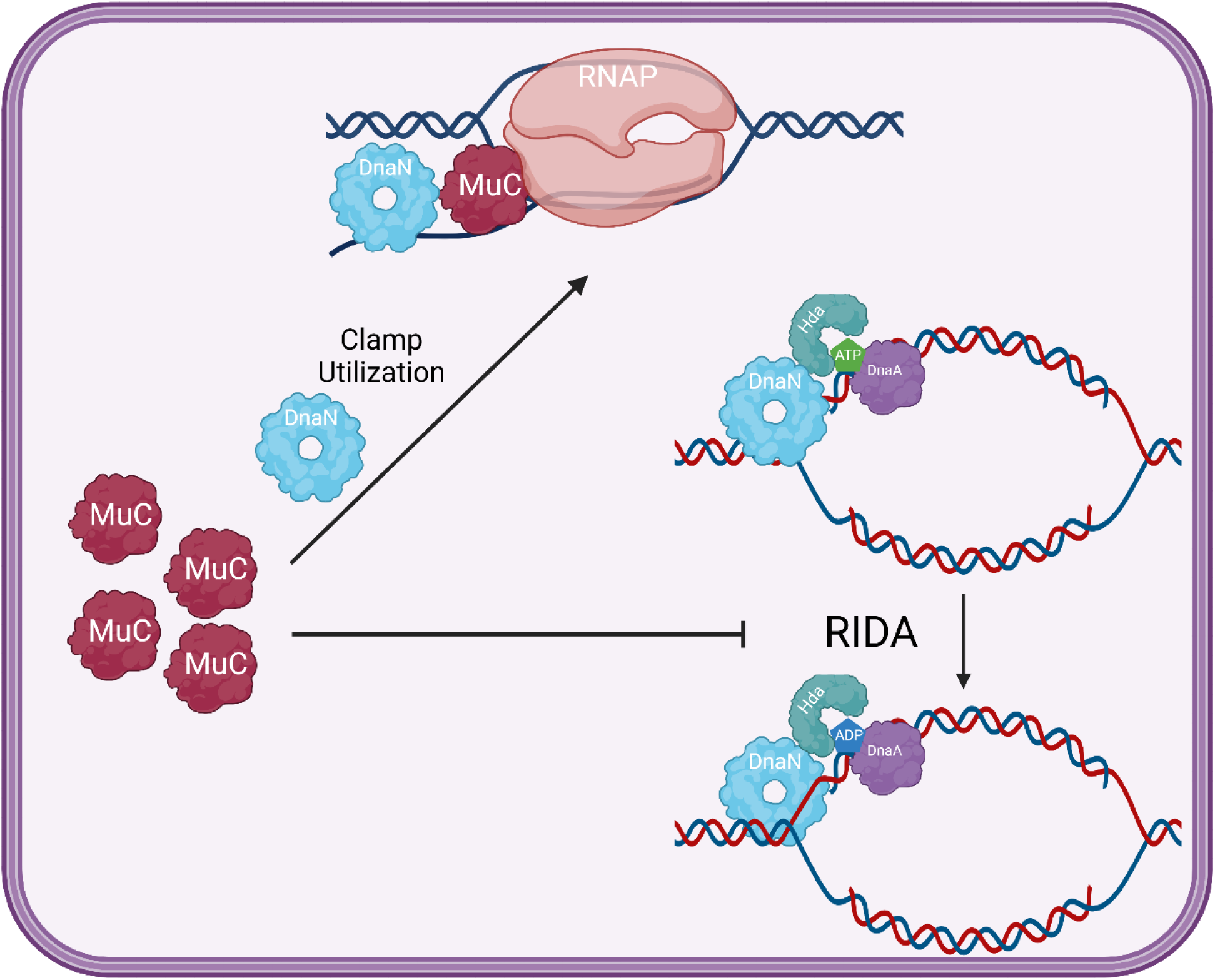
Model for MuC Toxicity. MuC, through its interaction with the Clamp-Binding Motif, depletes the available supply of cellular Clamp. This depletion of the clamp prevents proper regulatory inhibition of DnaA of the Clamp-Hda complex (the hydrolysis of ATP-DnaA to ADP-DnaA), leading to over-initiation of replication, and consequently, to toxicity.

The observation that overexpression of DnaA could suppress MuC transcription, and that mutants of DnaA that suppresses MuC toxicity effectively reverse this suppression (Fig. S1) was a surprising finding since DnaA is not known to be required for Mu development [47]. Our current explanation for this observation is that an increased amount of DnaA would result in an increase in RIDA complex formation, though not necessarily the number of replication initiation events. This would, in turn, prevent MuC from accessing the clamp, thus delaying late gene transcription. Alternatively, it is conceivable that unavailability of Pol III components for the replication restart pathway (needed for repair of the transposition product [60,61]) due to assembly of the pre-replication complex at *oriC*, irrespective of whether replication initiation occurs, is sufficient to prevent Mu transcription due to decreased Mu copy number. Both models are not mutually exclusive since they both rely on the basic premise that Pol III components are limited and differ only in which of these limited components of Pol III Mu requires for replication, transcription, or both.

Regardless of the exact mechanism(s) of DnaA repression of MuC transcription, DnaA mutants that arose out of selection for MuC resistance have eeither lost the ability to suppress MuC transcription or can stimulate it upon DnaA overexpression (Fig. S1B, Fig. S1C). Yet, these DnaA mutants are mostly proficient for all steps that leads to replication initiation (no copy of the Wt DnaA is available in these mutants), suggesting that a minor defect in one or more replication-related activities of DnaA is sufficient for a large effect on DnaA repression of MuC transcription. We do note that in the assay for repression of Mu transcription, a Wt allele of DnaA is available on the chromosome. This would enable formation of hybrid DnaA oligomers, and perhaps, it is only the hybrid oligomer that is alleviating the repression activity of DnaA. Regardless of the complexity of DnaA hybrid and considering that MuC transcription utilizes the clamp, we suggest that ATP-DnaA is not as available in these DnaA mutants, thus reducing the amount of RIDA complex available and enabling MuC transcription. How this would be achieved mechanistically would be an interesting question to resolve. We note that DnaA R342H has been previously isolated as a cold-sensitive variant [62]. This variant is defective for duplex opening but is otherwise indistinguishable from Wt for ATP binding and *oriC* binding [63]. Since the two DnaA mutants that arose from MuC selection are either a small deletion in the linker domain II, which has been shown to mediate the interaction between DnaA and acidic phospholipid [54], and a point mutation in domain III, near the amphipathic helix, which is thought to directly interacts with the membrane [64,65], we hypothesize that these DnaA mutants are altered for membrane interactions, causing a decrease in ATP-DnaA, which would relieve the toxicity of MuC overexpression.

Toxicity associated with proteins interacting with the clamp has been reported previously. Both the target recognition motif, TnsE, of Tn7 [66] and the initiation protein, TrfA, of the RK4 plasmid has been shown to bind to the clamp [67]. Overexpression of TnsE leads to induction of the SOS response, and this induction of response is dependent on the CBM [66]. Toxicity of TrfA overexpression, which is also dependent on the CBM, is suppressible by corresponding overexpression of *hda* [68], but it is not clear over-initiation is occurring in this system. Curiously, we found that MuC overexpression did not cause induction of the SOS response (Fig. S3C, Fig. S3D), even though the clamp interaction is absolutely required for toxicity (Fig. 3B). We suspect that this difference may reside in the fact that MuC binds to RNAP to facilitate transcription of Mu late genes and not proteins involved in targeting DNA replication like TrfA or Tn7. More specifically, we think that MuC bound to the clamp, in the absence of a promoter, does not have any activity (i.e. DNA binding or transcription activation), and thus, depleting the intracellular pool of available clamp without interfering with other components of the replication machinery. This model is based on the observation that Hda was identified as a multi-copy suppressor of TrfA [68], while Hda overexpression alone is insufficient to rescue MuC toxicity (Fig. S2E).

Even though MuC toxicity and MuC transcription both involve the clamp, mutants with comparable levels of MuC-clamp interaction produce differing results for these phenotypes. In the case of toxicity, only the L5A mutation abolishes toxicity, and the Q2A mutation has no effect (Fig. 3B, Fig. S3B). By contrast, Q2A mutation produces transcriptional activity at an intermediate level between Wt and L5A (Fig. 4A), despite having similar level of MuC-clamp interaction as the Wt (Fig. 3A). Furthermore, the Q2 residue is heavily conserved among Mor/MuC family of transcription, rendering the lack of phenotype upon mutation alanine substitution at this position puzzling. We speculate that this difference is attributable to the method by which transcription by MuC is being measured. *lacZ*, a 3kb gene, is longer than the average gene length (approximately 1kb), thus exacerbating the need for clamp-MuC interaction for productive transcription of the full-length mRNA. This would enable a mutation that would otherwise have little effect on MuC-clamp to produce a phenotype, and not representative of the importance of this position for MuC-clamp interaction. Alternatively, it is conceivable that the MuC-clamp level of the interaction that is required for transcription (whether it is initiation or elongation), is much higher than simple sequestration of the clamp from the Hda to produce the over-initiation phenotype. We favor the former model since Mu prophages carrying MuC mutations in the CBM do not exhibit severe defects in late gene transcription (Fig. 4D, Fig. S4B).

The utilization of the clamp for transcription by MuC is reminiscent of, and perhaps analogous to, the coupling of DNA replication to activation of late gene transcription of T4 phage [27]. This coupling of replication to activation late gene transcription requires a specific DNA structure, either a nick or primer/template junction, gp33/ gp55, both which are RNAP binding proteins, and gp45, the T4 clamp [27]. The protein component of this coupling system is analogous to that of MuC-clamp transcription: MuC could be thought of as an equivalent of the gp33/gp55 complex since MuC shares similarity with gp33/gp55 in that a -10 consensus is recognizable, but a -35 consensus is not evident. T4 clamp and the *E. coli* clamp are similar in that they both form a doughnut-like structure which encircles the DNA, though the T4 clamp is prone to opening in solution [69,70]. Given that T4 couple replication to late gene transcription through the availability of a nicked DNA or primer/template junction [27], we speculate that MuC-clamp interaction may also serve to couple late gene activation to replication, in addition to any other effect the clamp might have (e.g. processivity for transcription of a long template). We find this model to be attractive due to the well-understood product of Mu replication: Mu transposition results in a nick on the 3’-OH at the target site. Since Mu transposition has been well-described *in vitro*, and the products of transposition examined in detail, one should be able examine whether these products are competent for MuC-clamp transcription *in vitro*.

## Acknowledgments

This work was supported by NIH grant GM118085.

## Materials and Methods

### Media, Strains, Phages and Plasmids

Unless conditions are specified, all strains are grown in LB at 30°C with shaking. When appropriate, antibiotics were at the following concentrations: Ampicillin (Amp) at 100µg/mL, Kanamycin (Kan) at 25µg/mL, Chloramphenicol (Cam) at 20µg/mL, and Streptomycin (Strep) at 100µg/mL. Anhydrotetracycline (aTc) was used for induction of Tet promoter at 200ng/mL. Isopropyl-β-D-Galactoside (IPTG) was used at 1mM. 5-bromo-4-chloro-3-indolyl-β-D-galactopyranoside (X-gal) was used at 80µg/mL. o-nitrophenyl-β-D-Galactoside (ONPG) was purchased from Sigma. Electrocompetent cells for transformation were prepared by washing a growing culture of O.D. 0.4-0.5 in cold 10% glycerol 3 times. The pellet was resuspended in 1:50 of the original volume in 10% glycerol. Electroporation was performed in *E. coli* Pulser (Biorad) with 1mm Electroporation cuvette Plus (Fisher) at 1.8kV. Cells were recovered in SOC (LB supplemented with 10mM MgCl_2_, 10mM MgSO_4_, and 0.2% Glucose) for 1.5 hours at 30°C with shaking prior to plating on the appropriate selection. Chemical competent cells were prepared by washing a growing culture of O.D. 0.4-0.5 in cold (10% glycerol, 20mM CaCl_2_) twice. Cells were subjected to heat shock at 42°C for 1 minute, recovered in ice-cold SOC for 2 hours, and plated on appropriate selection plate. Mu phage was stored in Mu Buffer (50mM Tris-HCl pH 8.0, 100mM NaCl, 5mM CaCl_2_, 5mM MgCl_2_, and 0.1% gelatin). Strains and Plasmids employed in this study are listed in Table S1. Primers, purchased from Integrated DNA Technology (IDT), used in this study are listed in Table S2.

### General Molecular Techniques

Routine PCR were performed with Taq DNA Polymerase (NEB) or Phusion Master Mix (Thermo Fisher), according to manufacturer’s instructions. PCR fragments for cloning were generated with Phusion DNA Polymerase (NEB) according to manufacturer’s instructions. Gibson Master Mix (isothermal) was made according to [71]. Gibson assembly was performed at 50°C for at least 2 hours, but typically is performed overnight. Golden Gate Assembly was performed with Esp3I (NEB), T7 DNA Ligase (NEB), T4 DNA Ligase Buffer (NEB). Primers used to generate the gRNA were used at 10nM each. 100ng of the destination vector was used. The program for Golden Gate Assembly was 3 minutes at 37°C, 2 minutes at 16°C for 35 cycles, followed by 1 cycle of 10 minutes incubation at 37°C. T4 DNA Ligase, T4 PNK, and T4 DNA Ligase Buffer were purchased from NEB.

### Plasmid Construction

Plasmids were constructed using Gibson assembly. BW25113 was used for transformation of plasmids. Typically, 250µL of the backbone was assembled with a molar equivalent of insert in a 20µL reaction. The resulting product was purified with PCR clean-up kit (Qiagen) according to the manufacturer’s instructions. 2µL of the 20µL eluted product was used for transformation. Colonies were screen by PCR with primers spanning the junction between the backbone and the insert. Positive clones were restruck and checked once more using the same primer pairs. The PCR product was sequenced to confirm the identity of the sequence by either Eton Biosciences or Azenta. For generating point mutants, primers carrying the desired mutation were used to amplify the plasmid of interest. The resulting PCR product was gel-extracted with Qiagen Gel Extraction Kit. The product was then self-ligated overnight with T4 DNA Ligase and T4 DNA PNK in T4 Ligase Buffer. The ligated product was then purified and 2µL of the 20µL eluted product was used for transformation. Positive clones carrying the desired mutation were identified by PCR of the target sequence followed by sequencing. The positive clones were then restruck once again and verified with PCR and sequencing.

### Strain/ Prophage Construction

For construction of deletion strains, pKD4 was used as the template, and the procedure is based on that of [72]. Briefly, 0.5mL of an overnight culture of MG1655 carrying pKD46 was pelleted, washed twice in 1mL of PBS, and diluted 1:100 in fresh LB supplemented with 0.2% arabinose. The culture was grown to an O.D. of approximately 0.5. The cells were made electrocompetent. Cells were then transformed with the appropriate PCR product and let recover in SOC for 3 hours at 30°C. The outgrowth was then plated on the appropriate selection and incubated at 37°C overnight. Positive clones were identified by PCR with primers amplifying the junction of expected insertion. Positive clones were struck out on the appropriate selection plate at 37°C and reconfirmed with PCR followed by sequencing. To remove the antibiotic-encoding cassette, pCP20 was transformed into the desired host strain. Clones carrying pCP20 were then struck out on LB plate without selection and incubated at 42°C overnight. Colonies were then checked for the loss of both pCP20 and the antibiotics cassette by streaking on the appropriate selection.

Construction of Mu prophage with desired point mutants is based on that [73]. Briefly, lysogens transformed with pCR2.0 (containing Lambda Red proteins-Arabinose inducible, and Cas9-aTc inducible) are grown overnight in the appropriate antibiotics. 0.5mL of the overnight culture was pelleted, washed twice in 1mL of PBS, and diluted 1:100 in fresh LB supplemented with 0.2% arabinose. The cells were grown to an O.D. of 0.4-0.5, made competent by washing with ice-cold 10% glycerol as described in the previous paragraph, and resuspended in 1:50 volume of 10% glycerol. To the mixture, 200µg of pKH314, 200µg of pKH315, and 2µL of a 100µM repair oligo, which determine what mutations would be introduced, were added to the mixture and mixed. The cells were electroporated and recovered in SOC for 1 hour at 30°C. After 1 hour, Kan was added to a final concentration of 50µg/mL. We note that we did not add aTc for induction since we found that we did not need TetR repression (i.e. no extra plasmid carrying the Tet repressor) for efficient editing. The culture was then grown for another 2 hours at 30°C and then 1µL, 10µL, and 100µL of the culture was plated on plates containing Amp and Kan and grown at 30°C overnight. Colonies from the plate with the lowest number of colonies were PCR to check for positive clone for editing. Positive clones were then restruck on Amp plate and verified again with PCR. After which, colonies were grown in LB supplemented with 0.2% rhamnose to remove pKH314/pKH315.

### Flow Cytometry

The procedure is based on [41,74]. Briefly, strains to be examined were grown overnight in LB at 30°C in the appropriate antibiotic(s). 0.5mL of the washed twice in 1mL of PBS and resuspended in 0.5mL of PBS. For strains carrying MuC overexpression vector, the suspension was diluted 1:100 in 10mL of LB supplemented with aTc supplemented with Kan. Otherwise, cells were diluted 1:100 in 10mL of LB. The culture was grown to an O.D. of 0.2. Ciprofloxacin and Rifampicin was added to the culture to a final concentration of 10µg/mL and 120µg/mL, respectively. The cells were then continued to be incubated at 30°C for another 4 hours. The cells were then pelleted by centrifugation (4500g, 4°C, 10 minutes). The cells were fixed by the addition of 70% ethanol (50mL), incubated at room temperature for 10 minutes. The cells were then pelleted and washed with 50mL PBS twice. The pellet was then flash frozen for subsequent analysis the next day. The pellet was resuspended in 10mL of PBS and RNaseA (NEB) was added to final concentration of 100µg/mL and incubated at 37°C for 4 hours. The cells were then pelleted and resuspended in 10mL of PBS. For DNA labelling, the cell suspension was diluted 1:10 in 1mL of diH_2_O and Sytox Green (Molecular Probes?) was added 1:1000 v/v and incubated at 37°C for 30 minutes. The cells were then analyzed on LSRFortessa SORP Flow Cytometer (BD Biosciences) using filter settings for FITC. 10,000 cells were analyzed for each sample. Chromosomal equivalent was determined by using cells from an overnight culture subjected to fixation and subsequent staining. Three biological replicates were analyzed and one is shown.

### MuC Toxicity Assay

Strains to be examined were grown overnight in LB at 30°C in the appropriate antibiotic(s). 0.5mL of the washed twice in 1mL of PBS and resuspended in 0.5mL of PBS.

For plate-based assay, 10µL of serial dilution of the suspension (10^-1^ to 10^-6^) was spotted on LB plate supplemented with the appropriate antibiotics and inducer (aTc and/or IPTG when needed). The plates were incubated overnight at 30°C and imaged the following day (typically 16 hours). For colonies to form with MuC induction, plates were incubated further at 30°C for another 24 hours before colonies could be scored. The plate was imaged, and one of three biological replicates is shown.

For liquid growth, the suspension was diluted 1:100 in 10mL of LB supplemented with appropriate antibiotics and aTc, when MuC overexpression is desired. O.D. _600nm_ was monitored at indicated time points with Smart-Spec Plus spectrophotometer (Biorad). Measurement for three biological replicates were made. The mean and error are shown

### Isolation, Sequencing and Analysis of MuC Suppressor

BW25113 carrying MuC overexpression vector and p*mom*-*lacZ* was grown overnight in LB supplemented with Kan and Cam. 0.5mL of the overnight culture was washed twice in 1mL of PBS and resuspended in 0.5mL of PBS. 100µL of serial dilutions (10^-1^ to 10^-4^) of the suspension was plated on LB plate supplemented with Kan, Cam, aTc, and X-gal and incubated at 37°C overnight. Blue colonies were then struck out on LB plate supplemented with Kan, Cam, aTc, and X-gal again to confirm resistance to MuC and p*mom* activity. 14 positive clones were picked and grown in LB supplemented with Kan overnight. Plasmids were extracted and transformed into Wt BW25113 to confirm that MuC from these plasmids were still functional. Plasmids from positive clones were still toxic in Wt strain. Genomic DNA was prepared from these resistant strains along with the parent strain as control with Wizards Genomic DNA Kit (Promega) according to manufacturer’s instructions. Sequencing was performed by Novogene on the NovaSeq platform, paired end reads 150. Reads were mapped to the genome of BW25113 (GCA_000750555.1) using default parameters on Geneious. SNPs, indels and large rearrangement were detected with Geneious using default parameters. SNPs and indels present in the parent strain were discarded from analysis. SNPs and indels were scored as a hit if the variant frequency was greater than 75%. Mutants and the associated mutations are shown in Table 1.

### Mu Phage Preparation

When required, lysogen of the indicated prophage was grown overnight, diluted 1:100 in fresh LB and grown until an O.D. of 0.5. After which, the culture was shifted to a 42°C waterbath and incubated until lysis is complete. The lysate was clarified by centrifugation at 4500g for 10 minutes. Chloroform was added to the mixture (10% v/v) and shaken. The phases were separated by centrifugation at 4000g for 10 mins at 4°C. The aqueous phase was collected (top layer) and used to assay Mu plaquing.

### Mu Lysis

0.5mL of an overnight culture of the desired lysogen was grown in LB supplemented with the appropriate antibiotics was pelleted, washed twice in 1mL of PBS, and diluted 1:100 in 10mL of LB in a 50mL conical. The culture was grown to an O.D. _600nm_ of 0.3-0.4 at 30°C. The culture was then shifted to a 42°C waterbath and incubated with shaking. At indicated time points, 700µL was withdrawn, and O.D. _600nm_ was measured.

### Western Blot

BW25113 carrying vectors expressing MuC-FLAG or MuC variants were grown overnight in LB supplemented with Kan. .5mL of the overnight culture was washed twice in 1mL of PBS and resuspended in 0.5mL of PBS. The suspension was diluted 1:100 in 10mL of LB supplemented with Kan and grown to an O.D. of 0.5-0.6. MuC overexpression is non-toxic at this O.D. value. aTc was added to induce expression of MuC, and the cells were grown for an additional 2 hours. Cells were harvested by centrifugation (4500g, 4°C, 10 minutes), washed in 10mL of PBS twice, and flash frozen in liquid N_2_. The pellet was then resuspended in 2mL of PBS, and sonicated (Vibracells, Tip model CV33) on a 10s on-10s off cycle in the cold room for 1 minute total. The debris was removed via centrifugation (4500g, 4°C, 10 minutes). The supernatant was collected and used for subsequent steps. 10uL of the supernatant and 2µL of the loading was boiled at 98°C for 10 minutes. The mixture was loaded onto an SDS-PAGE gel (typically 10% polyacrylamide) and ran for 75 minutes or until the dye reaches the bottom at 150V in running buffer (25mM Tris, 0.192mM Glycine,0.1% SDS). Proteins were then transferred onto membrane (Protran BA85) at 100V for 1 in cold transfer buffer (25mM Tris, 0.192mM Glycine, 0.1% SDS, 20% Methanol). The membrane was then blocked in blocking buffer (Intercept Blocking Buffer, Licor) for 1 hour at room temperature. The membrane was then incubated with α-DnaK (Mouse, Enzo Lifesciences, ADISPA880F) and α-FLAG (Mouse, Sigma, F180450UG) in blocking buffer (10mL) overnight at 4°C. The membrane was then washed 3 times with 10mL of block buffer. The membrane was then probed with secondary antibody (α-Mouse IR680, Licor, NC0252290). The blot was then visualized on Odyssey Clx (Licor).

### Alignment of Mor/MuC family of transcription factors

BlastP was performed on MuC sequence [75]. Aligned sequences (100 total) were downloaded and aligned with ClustalW on Geneious. Sequence logo threshold for the consensus was set to 95%.

### Miller Assay

At indicated time points, 20µL of the indicated culture was withdrawn, its O.D. _600nm_ recorded on Smart-Spec Plus spectrophotometer (Biorad). and added to 80µL of permeabilization buffer (100mM NaHPO_4_, 20mM KCl, 2mM MgSO_4_, 0.8mg/mL CTAB, 0.4mg/mL sodium deoxycholate, 5.4µL/mL β-mercaptoethanol). After the final time point, 600µL of substrate solution (60mM Na_2_HPO_4_, 40mM NaH_2_PO_4_, 1mg/mL ONPG, 2.7µL/mL β-mercaptoethanol) was added to each sample. The samples were incubated at 30°C for variable times depending on source of MuC expression: for plasmid-borne MuC, incubation time is typically 20-30 minutes; for MuC from the Mu prophage, 1 hour is often necessary before appreciable signal could be observed. 600µL of 1M Na_2_CO_3_ was added to stop the reaction. O.D. _420nm_ was recorded on Smart-Spec Plus spectrophotometer (Biorad). Miller units were calculated as follows:

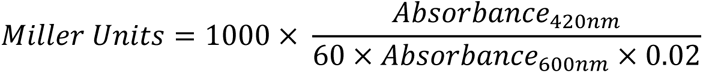

### Temperature Shift Experiments

MuC specific transcription was assayed with pKH33 (p*mom-lacZ*). σ^D^ transcription was assayed with pKH34 (p*Tet-lacZ*). Either of these plasmids were transformed into strains carrying the appropriate temperature sensitive allele strain carrying pKH32 and selected on LB plate supplemented with Kan and Cam. Single colonies were picked and grown overnight at 30°C (permissive temperature) supplemented with Kan and Cam. When indicated, strains were also transformed with plasmid carrying Wt DnaN (pKH45) and Amp was added to all subsequent selection step. 0.5mL of the culture was washed twice in 1mL of PBS, resuspended in 0.5mL of PBS, and diluted 1:100 in 20mL of LB (10mL each in 2 50mL conicals) with the appropriate antibiotics. The culture was grown an O.D. of 0.2 and the 2 conical was combined and re-split into 2 50mL conical (10mL each). One conical was incubated at 30°C the other was shifted to 42°C (non-permissive temperature). Miller assay was carried out as described above at the indicated time points. Relative transcriptional activity is defined as the Miller units of the culture at non-permissive to that of permissive temperature.

### Bacterial Two Hybrid Assay

The host strain for bacterial two-hybrid, BTH101, was transformed with the indicated plasmids and selected on plates supplemented with Amp, Kan, and Strep. Single colonies were picked and grown overnight in LB supplemented Amp Kan, and Strep. 0.5mL of the culture was washed twice in 1mL of PBS, resuspended in 0.5mL of PBS. 10µL of the washed cells were plated on LB plates supplemented Amp, Kan, Strep, and Xgal. The plates were incubated at 30°C for 16-24 hours, since color-development time seems to be variable. For quantification of β-galactosidase activity, cells on the plate were scrapped off and diluted to an O.D. between 0.4-0.6 in PBS. Miller assay was performed on this dilution as described above.

## Supplementary Figures and Tables

**Figure S1.**
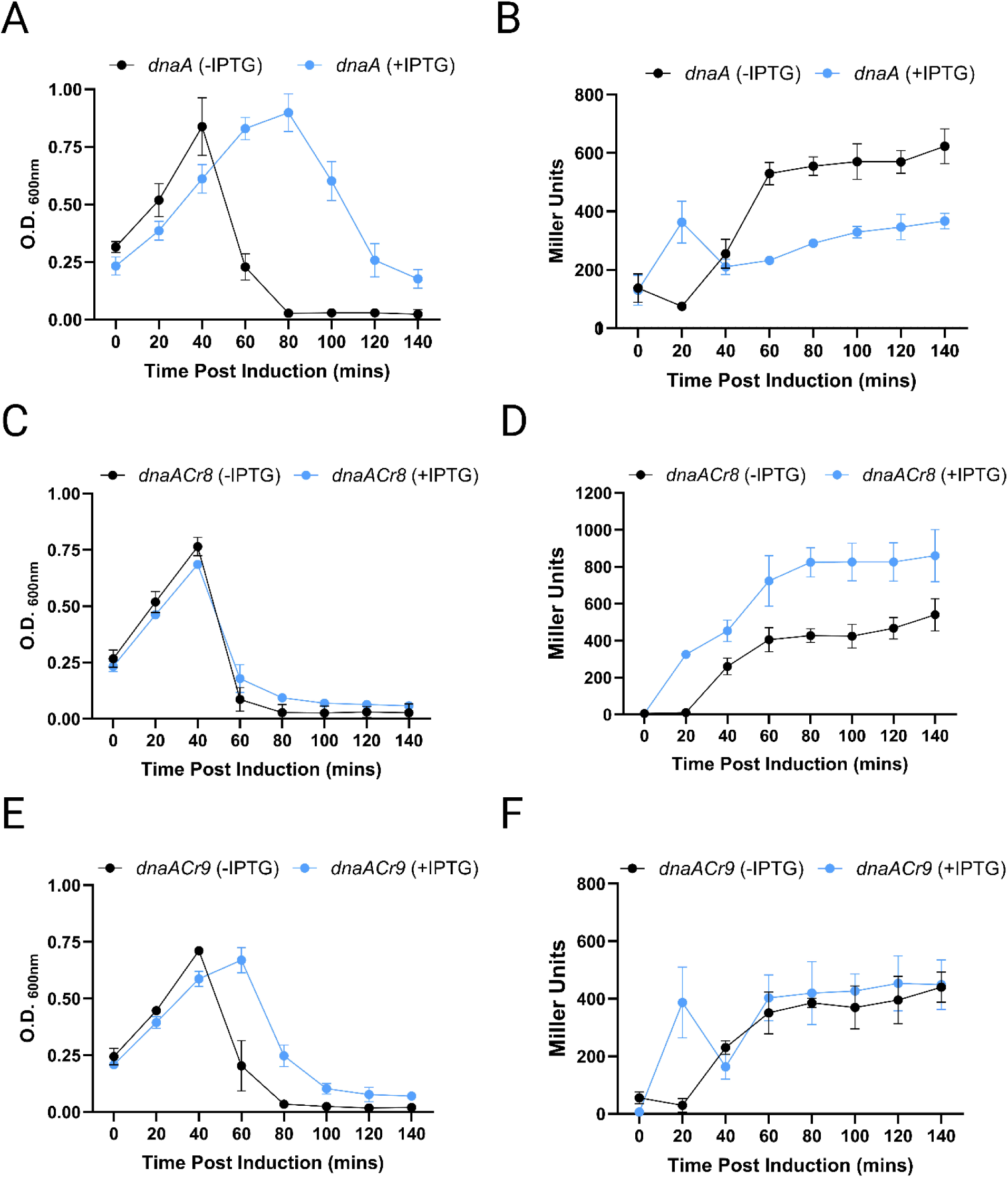
Lysis curves and late gene transcription of Mu lysogens supplied with DnaA variants. A, C, and E. Lysis curves for Mu lysogens carrying wt DnaA, DnaA Δ96-118, and DnaA R342H, respectively. B, D, F, Late gene transcription of the Mu lysogens from p*mom-lacZ* carrying wt DnaA, DnaA Δ96-118, and DnaA R342H, respectively.

**Figure S2.**
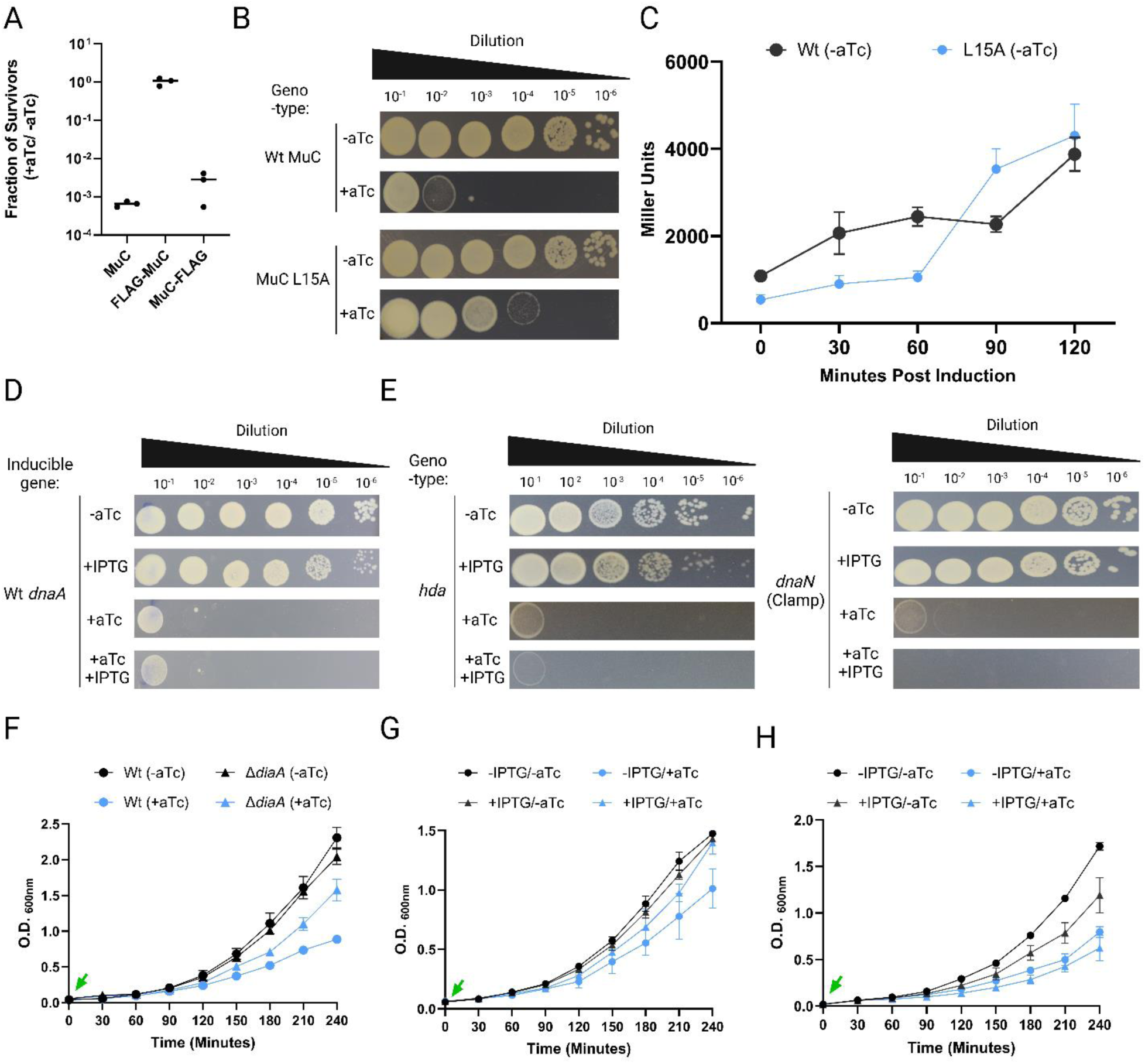
Rescue of MuC toxicity. A) Effect of FLAG fusion on MuC toxicity. B) MuC L15A is less toxic than Wt. C) MuC L15A is not defective for transcription. D) Wt DnaA is unable to rescue MuC toxicity. E) Hda and DnaN individually do not rescue MuC toxicity. F, G, and H. Rescue of MuC toxicity in liquid for Δ*diaA* background, expression of *dnaA* L366K, and expression of *hda-dnaN*, respectively. Green arrow indicates time of aTc addition.

**Figure S3.**
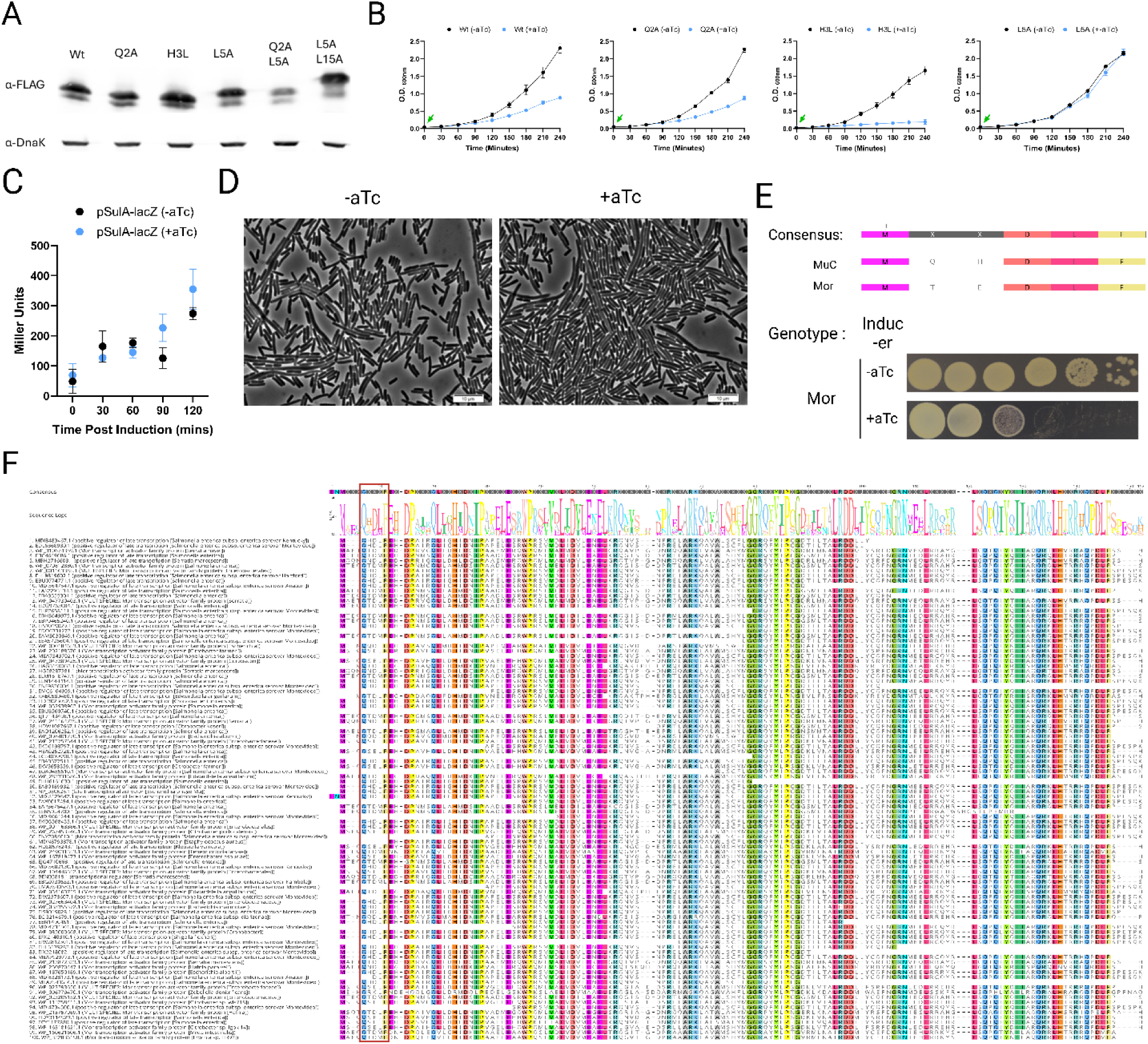
Effect of MuC-Clamp interaction. A) Western Blot of MuC variants in the Clamp Binding Motif (CBM). B) Effect of CBM mutations on MuC toxicity. C) Transcription of pSulA with MuC overexpression. D) Cell morphology with MuC overexpression. E) Toxicity of Mor overexpression. F) Alignment of 100 members of Mor/MuC family of transcription factors. Consensus sequence (95% threshold) is shown on top. Red box indicates the putative CBM.

**Figure S4.**
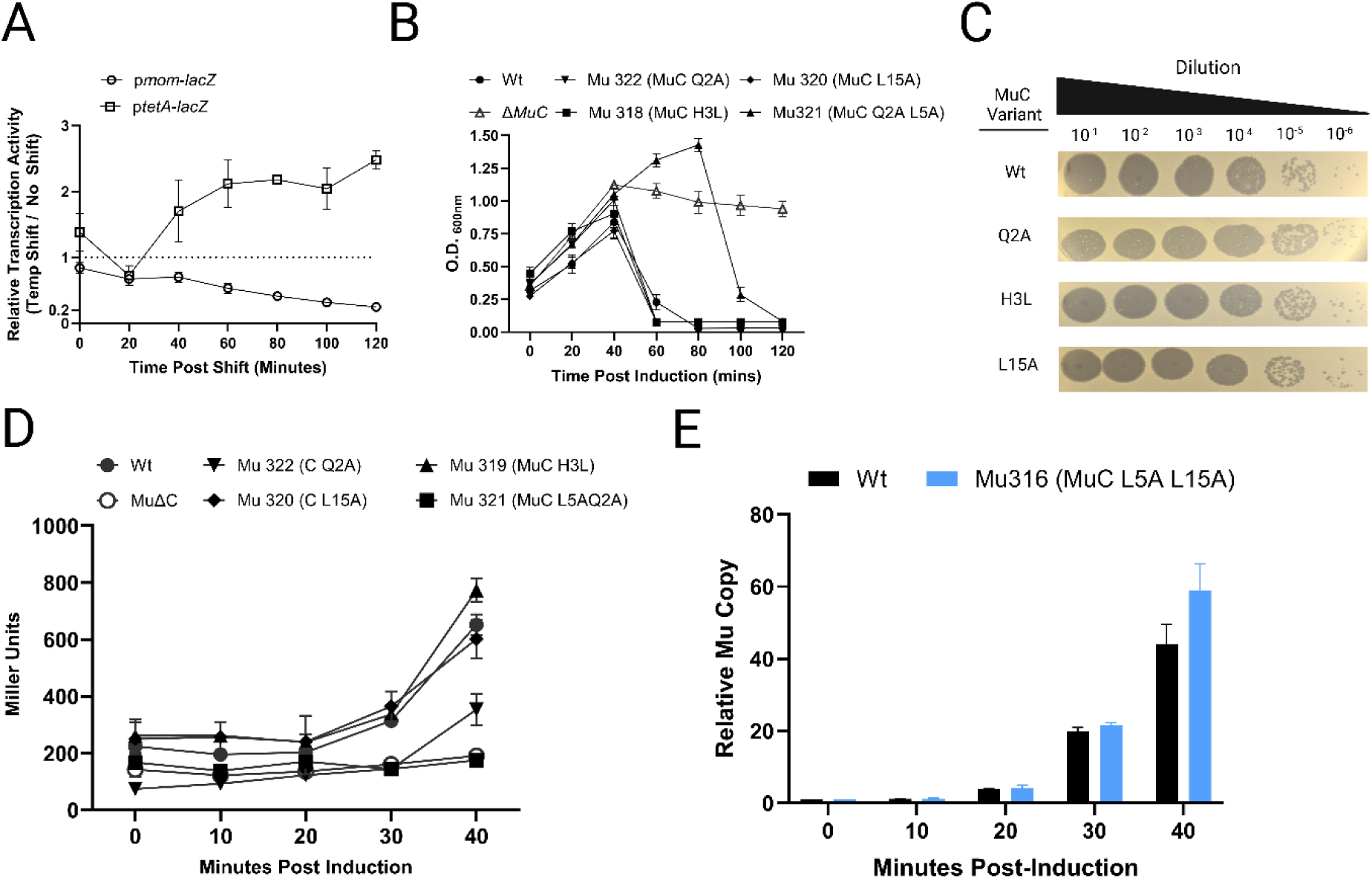
Dependence of Mu late gene transcription on the Clamp. A) Dependence of MuC transcription on the clamp loader (*dnaX)*. B) Lysis curve of Mu prophages carrying MuC variants that showed various levels of Clamp interaction. C) Plaquing of morphology prophages carrying MuC variants that showed various levels of Clamp interaction. D) Late gene transcription of prophages carrying MuC variants that showed various levels of Clamp interaction. E) Replication of Mu prophage defective for clamp interaction, lysis and transcription as quantified by qPCR.

**Table S1.**
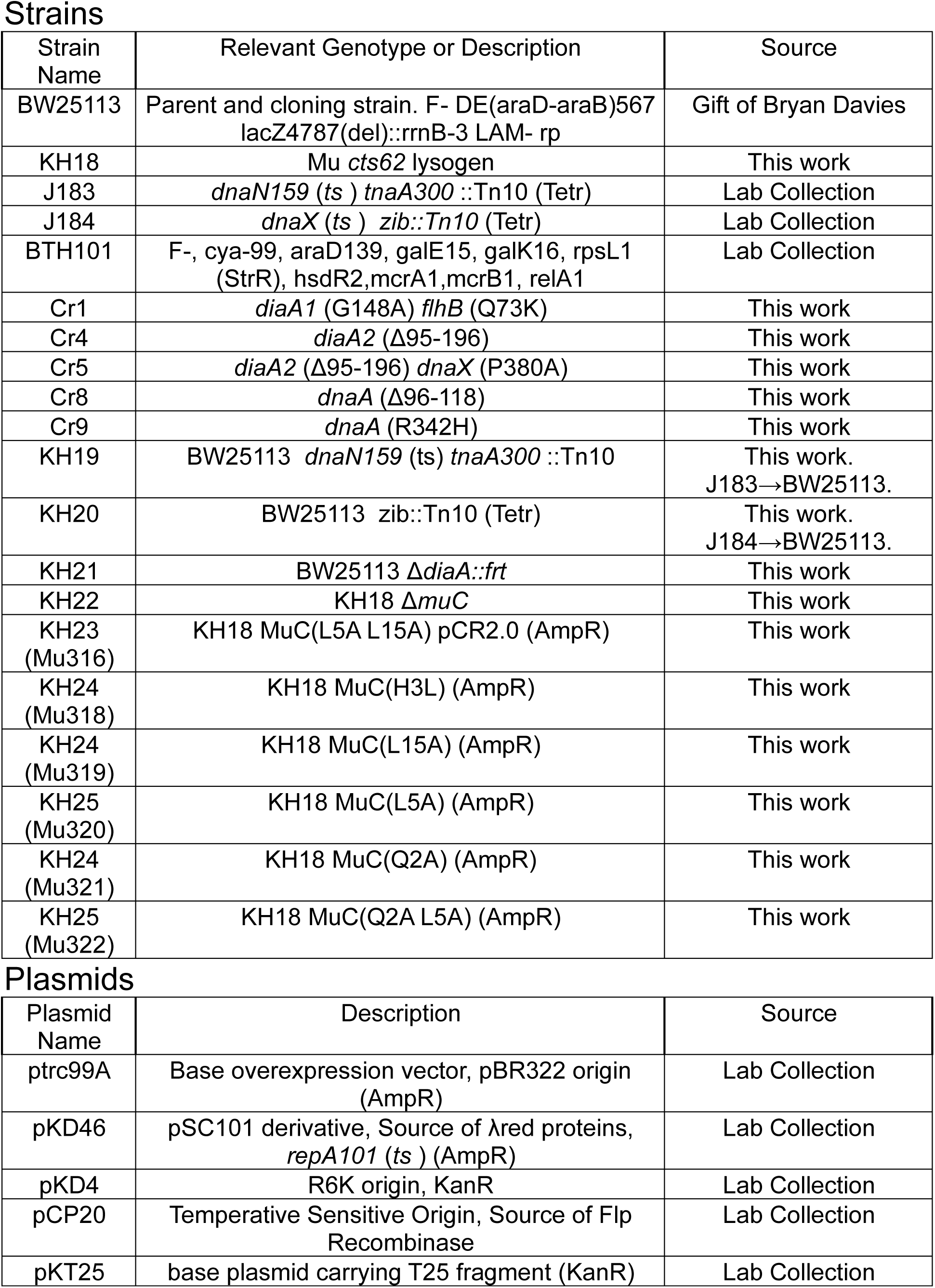
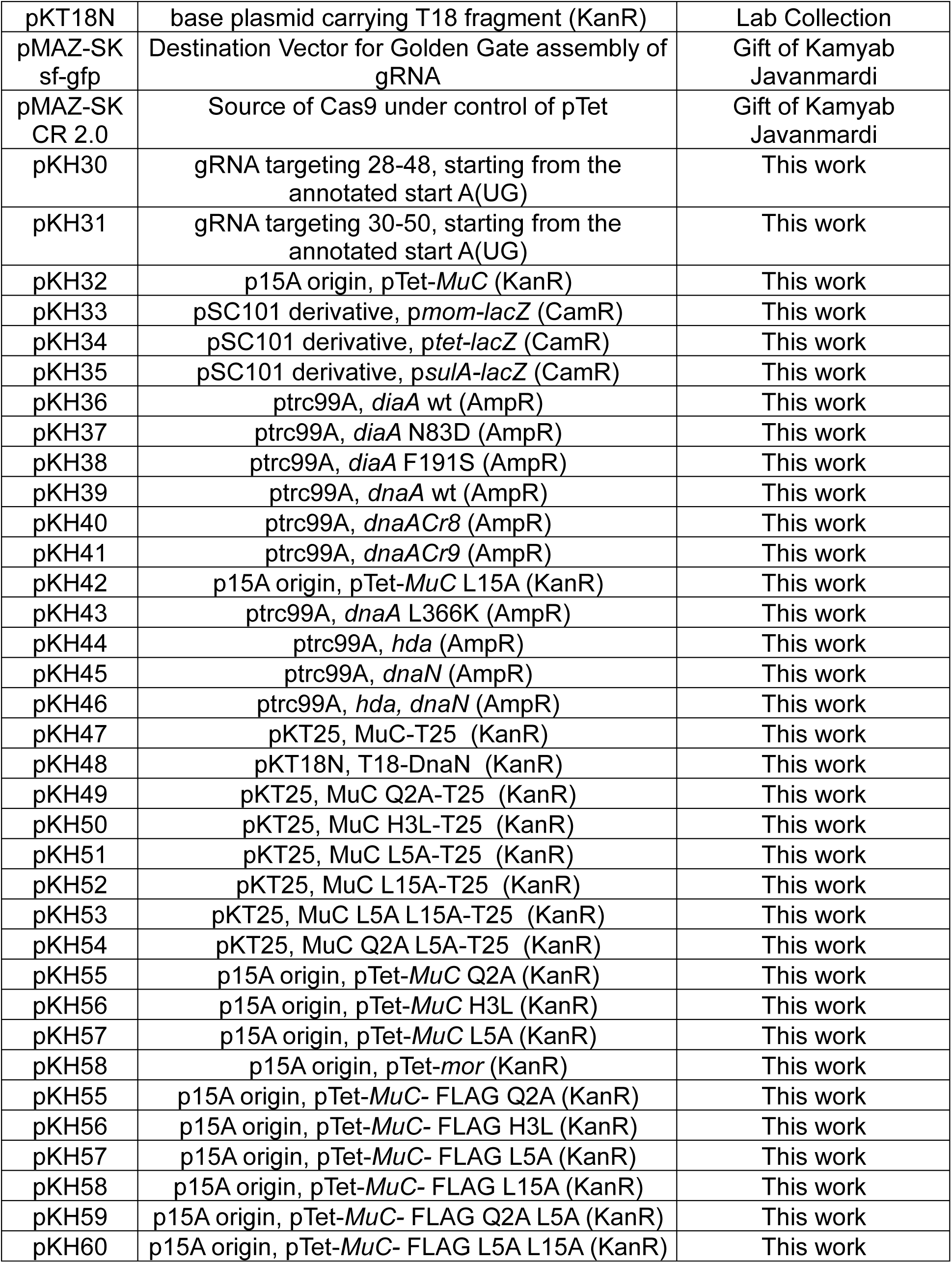
Strains and Plasmids used in this study.

Strains
| Strain Name | Relevant Genotype or Description | Source |
| --- | --- | --- |
| BW25113 | Parent and cloning strain. F- DE(araD-araB)567 lacZ4787(del)::rrnB-3 LAM- rp | Gift of Bryan Davies |
| KH18 | Mu *cts62* lysogen | This work |
| J183 | *dnaN159 (ts ) tnaA300 ::Tn10* (Tetr) | Lab Collection |
| J184 | *dnaX (ts ) zib::Tn10* (Tetr) | Lab Collection |
| BTH101 | F-, *cya-99*, *araD139*, *galE15*, *galK16*, *rpsL1* (StrR), *hsdR2*, *mcrA1*, *mcrB1*, *relA1* | Lab Collection |
| Cr1 | *diaA1* (G148A) *flhB* (Q73K) | This work |
| Cr4 | *diaA2* (Δ95-196) | This work |
| Cr5 | *diaA2* (Δ95-196) *dnaX* (P380A) | This work |
| Cr8 | *dnaA* (Δ96-118) | This work |
| Cr9 | *dnaA* (R342H) | This work |
| KH19 | BW25113 *dnaN159* (ts) *tnaA300 ::Tn10* | This work.<br>J183→BW25113. |
| KH20 | BW25113 *zib::Tn10* (Tetr) | This work.<br>J184→BW25113. |
| KH21 | BW25113 Δ*diaA::frt* | This work |
| KH22 | KH18 Δ*muC* | This work |
| KH23<br>(Mu316) | KH18 MuC(L5A L15A) pCR2.0 (AmpR) | This work |
| KH24<br>(Mu318) | KH18 MuC(H3L) (AmpR) | This work |
| KH24<br>(Mu319) | KH18 MuC(L15A) (AmpR) | This work |
| KH25<br>(Mu320) | KH18 MuC(L5A) (AmpR) | This work |
| KH24<br>(Mu321) | KH18 MuC(Q2A) (AmpR) | This work |
| KH25<br>(Mu322) | KH18 MuC(Q2A L5A) (AmpR) | This work |

Table S1. Strains and Plasmids used in this study.
| Plasmid Name | Description | Source |
| --- | --- | --- |
| ptrc99A | Base overexpression vector, pBR322 origin (AmpR) | Lab Collection |
| pKD46 | pSC101 derivative, Source of λred proteins, *repA101 (ts )* (AmpR) | Lab Collection |
| pKD4 | R6K origin, KanR | Lab Collection |
| pCP20 | Temperative Sensitive Origin, Source of Flp Recombinase | Lab Collection |
| pKT25 | base plasmid carrying T25 fragment (KanR) | Lab Collection |
| pKT18N | base plasmid carrying T18 fragment (KanR) | Lab Collection |
| pMAZ-SK<br>sf-gfp | Destination Vector for Golden Gate assembly of<br>gRNA | Gift of Kamyab<br>Javanmardi |
| pMAZ-SK<br>CR 2.0 | Source of Cas9 under control of pTet | Gift of Kamyab<br>Javanmardi |
| pKH30 | gRNA targeting 28-48, starting from the<br>annotated start A(UG) | This work |
| pKH31 | gRNA targeting 30-50, starting from the<br>annotated start A(UG) | This work |
| pKH32 | p15A origin, pTet-*Mu*C (KanR) | This work |
| pKH33 | pSC101 derivative, *pmom-lacZ* (CamR) | This work |
| pKH34 | pSC101 derivative, *ptet-lacZ* (CamR) | This work |
| pKH35 | pSC101 derivative, *psula-lacZ* (CamR) | This work |
| pKH36 | *ptrc99A*, *diaA* wt (AmpR) | This work |
| pKH37 | *ptrc99A*, *diaA* N83D (AmpR) | This work |
| pKH38 | *ptrc99A*, *diaA* F191S (AmpR) | This work |
| pKH39 | *ptrc99A*, *dnaA* wt (AmpR) | This work |
| pKH40 | *ptrc99A*, *dnaACr8* (AmpR) | This work |
| pKH41 | *ptrc99A*, *dnaACr9* (AmpR) | This work |
| pKH42 | p15A origin, pTet-*Mu*C L15A (KanR) | This work |
| pKH43 | *ptrc99A*, *dnaA* L366K (AmpR) | This work |
| pKH44 | *ptrc99A*, *hda* (AmpR) | This work |
| pKH45 | *ptrc99A*, *dnaN* (AmpR) | This work |
| pKH46 | *ptrc99A*, *hda*, *dnaN* (AmpR) | This work |
| pKH47 | pKT25, *Mu*C-T25 (KanR) | This work |
| pKH48 | pKT18N, T18-DnaN (KanR) | This work |
| pKH49 | pKT25, *Mu*C Q2A-T25 (KanR) | This work |
| pKH50 | pKT25, *Mu*C H3L-T25 (KanR) | This work |
| pKH51 | pKT25, *Mu*C L5A-T25 (KanR) | This work |
| pKH52 | pKT25, *Mu*C L15A-T25 (KanR) | This work |
| pKH53 | pKT25, *Mu*C L5A L15A-T25 (KanR) | This work |
| pKH54 | pKT25, *Mu*C Q2A L5A-T25 (KanR) | This work |
| pKH55 | p15A origin, pTet-*Mu*C Q2A (KanR) | This work |
| pKH56 | p15A origin, pTet-*Mu*C H3L (KanR) | This work |
| pKH57 | p15A origin, pTet-*Mu*C L5A (KanR) | This work |
| pKH58 | p15A origin, pTet-*mor* (KanR) | This work |
| pKH55 | p15A origin, pTet-*Mu*C- FLAG Q2A (KanR) | This work |
| pKH56 | p15A origin, pTet-*Mu*C- FLAG H3L (KanR) | This work |
| pKH57 | p15A origin, pTet-*Mu*C- FLAG L5A (KanR) | This work |
| pKH58 | p15A origin, pTet-*Mu*C- FLAG L15A (KanR) | This work |
| pKH59 | p15A origin, pTet-*Mu*C- FLAG Q2A L5A (KanR) | This work |
| pKH60 | p15A origin, pTet-*Mu*C- FLAG L5A L15A (KanR) | This work |

**Table S2.**
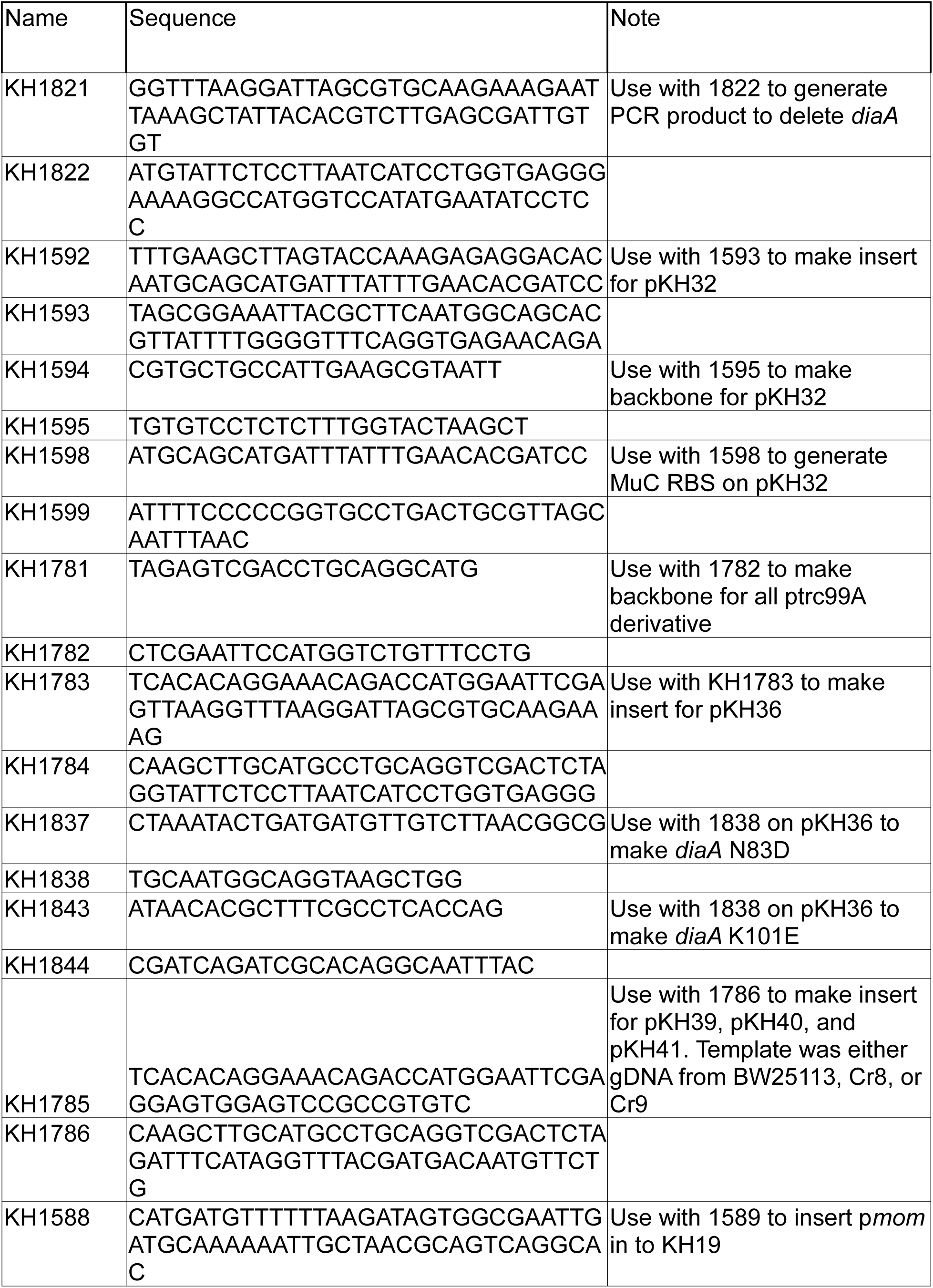
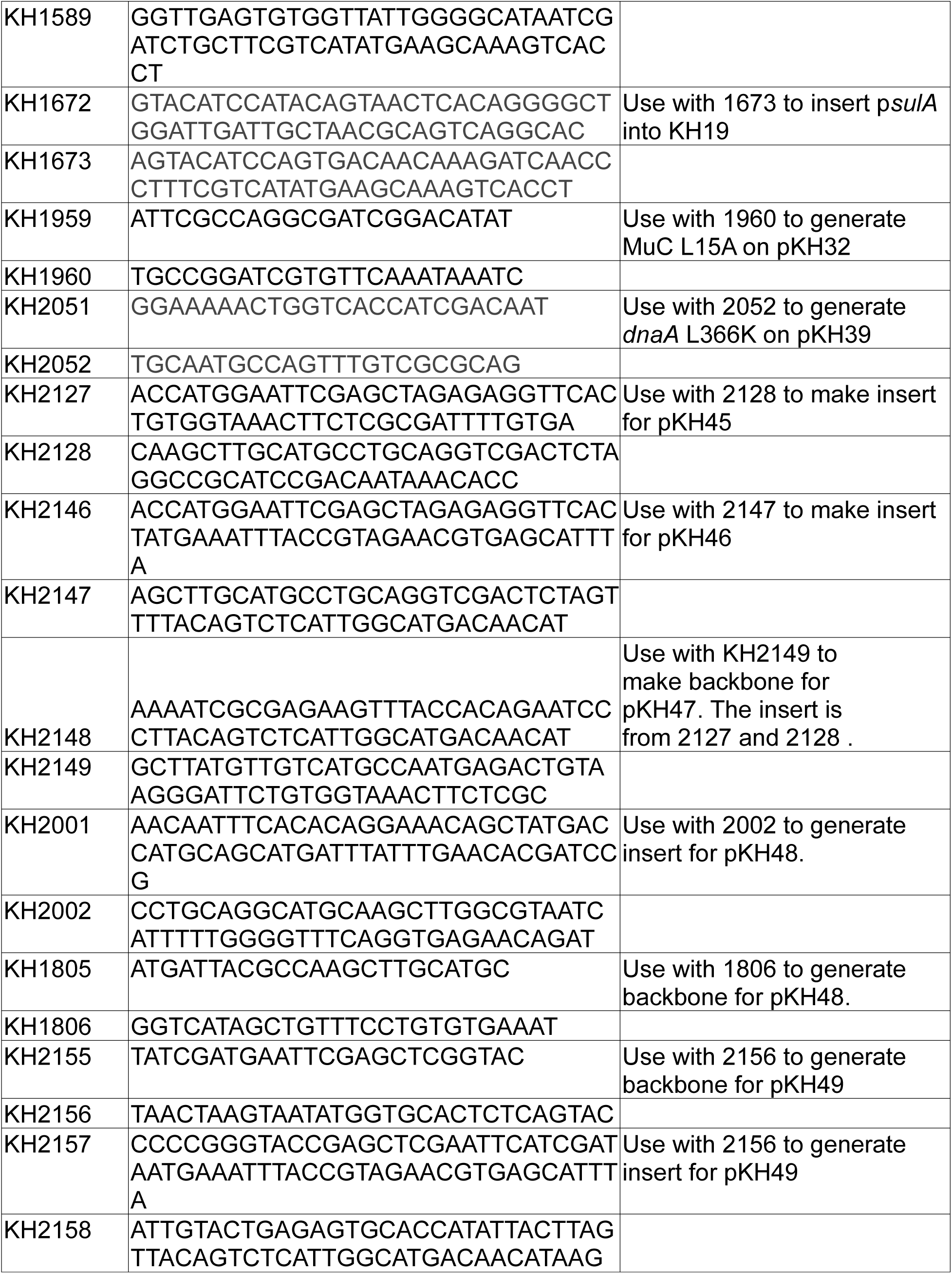
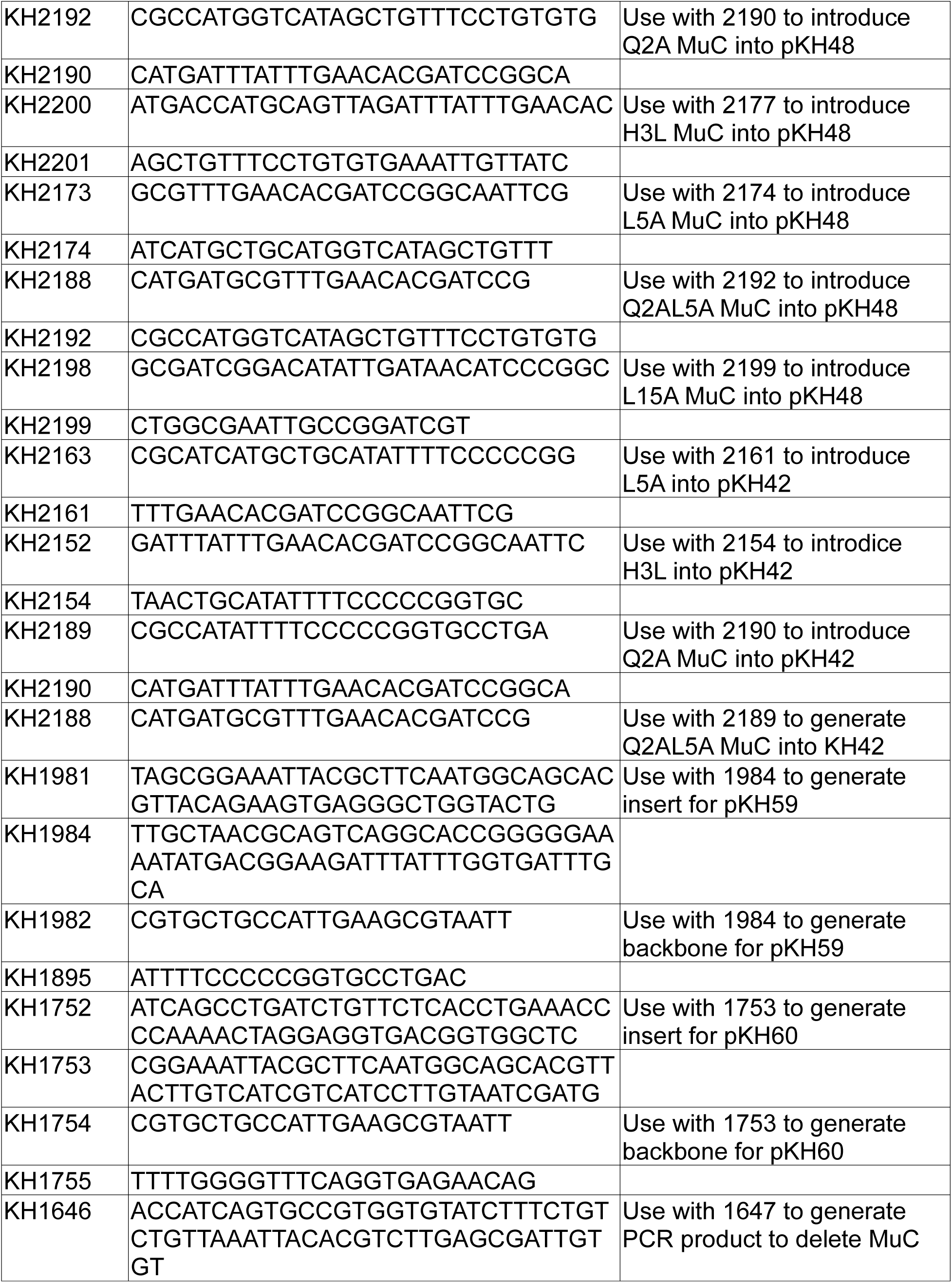
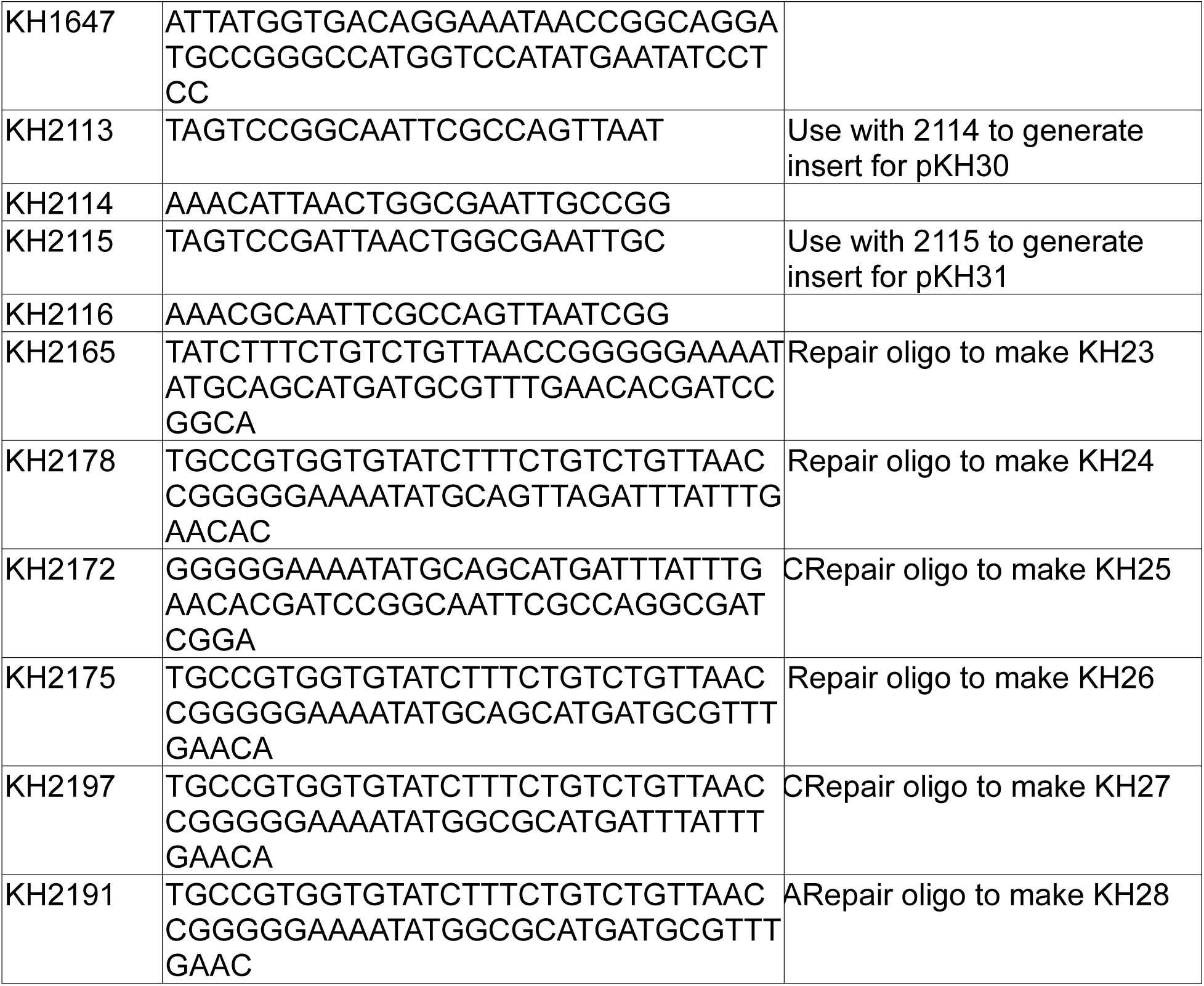
Primers Used in this study.

## References

1. M HR. Transposable Phage Mu. Microbiol Spectr. 2014;2: 10.1128/microbiolspec.mdna3-0007–2014. doi:10.1128/microbiolspec.mdna3-0007-2014

2. Chaconas G, Harshey RM. Transposition of Phage Mu DNA. Mobile DNA II. 2007. pp. 384–402. 10.1128/9781555817954.ch17

3. Mizuno N, Dramićanin M, Mizuuchi M, Adam J, Wang Y, Han Y-W, et al. MuB is an AAA+ ATPase that forms helical filaments to control target selection for DNA transposition. Proceedings of the National Academy of Sciences. 2013;110: E2441–E2450. doi:10.1073/pnas.1309499110

4. Zhao X, Gao Y, Gong Q, Zhang K, Li S. Elucidating the Architectural dynamics of MuB filaments in bacteriophage Mu DNA transposition. Nat Commun. 2024;15: 6445. doi:10.1038/s41467-024-50722-1

5. Mizuuchi K. Transpositional recombination: mechanistic insights from studies of mu and other elements. Annu Rev Biochem. 1992; 1011–51.

6. Nakai H, Doseeva V, Jones JM. Handoff from recombinase to replisome: Insights from transposition. Proceedings of the National Academy of Sciences. 2001;98: 8247–8254. doi:10.1073/pnas.111007898

7. Kelman Z, O’Donnell M. DNA polymerase III holoenzyme: structure and function of a chromosomal replicating machine. Annu Rev Biochem. 1995; 171–200.

8. Okazaki R, Arisawa M, Sugino A. Slow joining of newly replicated DNA chains in DNA polymerase I-deficient Escherichia coli mutants. Proceedings of the National Academy of Sciences 1972; 2954–2957. Available: 10.1073/pnas.68.12.2954

9. Fijalkowska IJ, Schaaper RM, Jonczyk P. DNA replication fidelity in Escherichia coli: a multi-DNA polymerase affair. FEMS Microbiol Rev. 2012;36: 1105–1121. doi:10.1111/j.1574-6976.2012.00338.x

10. Johnson A, O’Donnell M. Cellular DNA replicases: components and dynamics at the replication fork. Annu Rev Biochem. 2005; 283–315. Available: 10.1146/annurev.biochem.73.011303.073859

11. Maki H, Kornberg A. The polymerase subunit of DNA polymerase III of Escherichia coli. II. Purification of the alpha subunit, devoid of nuclease activities. Journal of Biological Chemistry. 1985;260: 12987–12992. 10.1016/S0021-9258(17)38825-7

12. Kong X-P, Onrust R, O’Donnell M, Kuriyan J. Three-dimensional structure of the β subunit of E. coli DNA polymerase III holoenzyme: A sliding DNA clamp. Cell. 1992;69: 425–437. 10.1016/0092-8674(92)90445-I

13. Pritchard AE, Dallmann HG, Glover BP, McHenry CS. A novel assembly mechanism for the DNA polymerase III holoenzyme DnaX complex: association of δδ’ with DnaX_4_ forms DnaX_3_δδ’; EMBO J. 2000;19: 6536–6545–6545. 10.1093/emboj/19.23.6536

14. Onrust R, Finkelstein J, Naktinis V, Turner J, Fang L, O’Donnell M. Assembly of a Chromosomal Replication Machine: Two DNA Polymerases, a Clamp Loader, and Sliding Clamps in One Holoenzyme Particle. I. ORGANIZATION OF THE CLAMP LOADER. Journal of Biological Chemistry. 1995;270: 13348–13357. 10.1074/jbc.270.22.13348

15. Xu Z-Q, Jergic S, Lo ATY, Pradhan AC, Brown SHJ, Bouwer JC, et al. Structural characterisation of the complete cycle of sliding clamp loading in Escherichia coli. Nat Commun. 2024;15: 8372. doi:10.1038/s41467-024-52623-9

16. Jeruzalmi D, Yurieva O, Zhao Y, Young M, Stewart J, Hingorani M, et al. Mechanism of Processivity Clamp Opening by the Delta Subunit Wrench of the Clamp Loader Complex of E. coli DNA Polymerase III. Cell. 2001;106: 417–428. 10.1016/S0092-8674(01)00462-7

17. Newcomb ESP, Douma LG, Morris LA, Bloom LB. The Escherichia coli clamp loader rapidly remodels SSB on DNA to load clamps. Nucleic Acids Res. 2022;50: 12872–12884. doi:10.1093/nar/gkac1169

18. Katayama T, Ozaki S, Keyamura K, Fujimitsu K. Regulation of the replication cycle: conserved and diverse regulatory systems for DnaA and oriC. Nat Rev Microbiol. 2010;8: 163–170. doi:10.1038/nrmicro2314

19. Mott ML, Berger JM. DNA replication initiation: mechanisms and regulation in bacteria. Nat Rev Microbiol. 2007;5: 343–354. doi:10.1038/nrmicro1640

20. Katayama T, Kubota T, Kurokawa K, Crooke E, Sekimizu K. The Initiator Function of DnaA Protein Is Negatively Regulated by the Sliding Clamp of the E. coli Chromosomal Replicase. Cell. 1998;94: 61–71. 10.1016/S0092-8674(00)81222-2

21. Kato J, Katayama T. Hda, a novel DnaA-related protein, regulates the replication cycle in *Escherichia coli*. EMBO J. 2001;20: 4253–4262–4262. 10.1093/emboj/20.15.4253

22. Kurokawa K, Nishida S, Emoto A, Sekimizu K, Katayama T. Replication cycle-coordinated change of the adenine nucleotide-bound forms of DnaA protein in *Escherichia coli*. EMBO J. 1999;18: 6642–6652–6652. 10.1093/emboj/18.23.6642

23. Su’etsugu M, Shimuta T, Ishida T, Kawakami H, Katayama T. Protein Associations in DnaA-ATP Hydrolysis Mediated by the Hda-Replicase Clamp Complex. Journal of Biological Chemistry. 2005;280: 6528–6536. 10.1074/jbc.M412060200

24. Miyachi K, Fritzler MJ, Tan EM. Autoantibody to a Nuclear Antigen in Proliferating Cells1. The Journal of Immunology. 1978;121: 2228–2234. doi:10.4049/jimmunol.121.6.2228

25. Taylor MS, LaCava J, Mita P, Molloy KR, Huang CRL, Li D, et al. Affinity Proteomics Reveals Human Host Factors Implicated in Discrete Stages of LINE-1 Retrotransposition. Cell. 2013;155: 1034–1048. doi:10.1016/j.cell.2013.10.021

26. Parks AR, Li Z, Shi Q, Owens RM, Jin MM, Peters JE. Transposition into Replicating DNA Occurs through Interaction with the Processivity Factor. Cell. 2009;138: 685–695. 10.1016/j.cell.2009.06.011

27. Geiduschek EP, Kassavetis GA. Transcription of the T4 late genes. Virol J. 2010;7: 288. doi:10.1186/1743-422X-7-288

28. Kassavetis GA, Elliott T, Rabussay DP, Geiduschek EP. Initiation of transcription at phage T4 late promoters with purified RNA polymerase. Cell. 1983;33: 887– 897. 10.1016/0092-8674(83)90031-4

29. Nechaev S, Kamali-Moghaddam M, André E, Léonetti J-P, Geiduschek EP. The bacteriophage T4 late-transcription coactivator gp33 binds the flap domain of Escherichia coli RNA polymerase. Proceedings of the National Academy of Sciences. 2004;101: 17365–17370. doi:10.1073/pnas.0408028101

30. Moarefi I, Jeruzalmi D, Turner J, O’Donnell M, Kuriyan J. Crystal structure of the DNA polymerase processivity factor of T4 bacteriophage11Edited by I. A. Wilson. J Mol Biol. 2000;296: 1215–1223. 10.1006/jmbi.1999.3511

31. Sanders GM, Kassavetis GA, Geiduschek EP. Use of a macromolecular crowding agent to dissect interactions and define functions in transcriptional activation by a DNA-tracking protein: bacteriophage T4 gene 45 protein and late transcription. Proceedings of the National Academy of Sciences. 1994;91: 7703–7707. doi:10.1073/pnas.91.16.7703

32. Wong K, Geiduschek EP. Activator-sigma interaction: a hydrophobic segment mediates the interaction of a sigma family promoter recognition protein with a sliding clamp transcription activator11Edited by R. Ebright. J Mol Biol. 1998;284: 195–203. 10.1006/jmbi.1998.2166

33. Howe M. Phage Mu. Symonds N, Toussaint A, Van de Putte P, Howe M, editors. Plainview, NY: Cold Spring Harbor Laboratory Press; 1987.

34. Bölker M, Wulczyn FG, Kahmann R. Role of bacteriophage Mu C protein in activation of the mom gene promoter. J Bacteriol. 1989;171: 2019–2027. doi:10.1128/jb.171.4.2019-2027.1989

35. Margolin W, Rao G, Howe MM. Bacteriophage Mu late promoters: four late transcripts initiate near a conserved sequence. J Bacteriol. 1989;171: 2003–2018. doi:10.1128/jb.171.4.2003-2018.1989

36. Margolin W, Howe MM. Activation of the bacteriophage Mu lys promoter by Mu C protein requires the sigma 70 subunit of Escherichia coli RNA polymerase. J Bacteriol. 1990;172: 1424–1429. doi:10.1128/jb.172.3.1424-1429.1990

37. Sun W, Hattman S, Kool E. Interaction of the bacteriophage mu transcriptional activator protein, C, with its target site in the mom promoter11Edited by R. Ebright. J Mol Biol. 1997;273: 765–774. 10.1006/jmbi.1997.1349

38. Weiyong S, Stanley H, Noboyuki F, Akira I. Activation of Bacteriophage Mu momTranscription by C Protein Does Not Require Specific Interaction with the Carboxyl-Terminal Region of the α or σ^70^Subunit of Escherichia coli RNA Polymerase. J Bacteriol. 1998;180: 3257–3259. doi:10.1128/jb.180.12.3257-3259.1998

39. Jiang Y, Howe MM. Regional mutagenesis of the gene encoding the phage Mu late gene activator C identifies two separate regions important for DNA binding. Nucleic Acids Res. 2008;36: 6396–6405. doi:10.1093/nar/gkn639

40. Messer W. The bacterial replication initiator DnaA. DnaA and oriC, the bacterial mode to initiate DNA replication. FEMS Microbiol Rev. 2002;26: 355–374. doi:10.1111/j.1574-6976.2002.tb00620.x

41. Ishida T, Akimitsu N, Kashioka T, Hatano M, Kubota T, Ogata Y, et al. DiaA, a Novel DnaA-binding Protein, Ensures the Timely Initiation of Escherichia coli Chromosome Replication. Journal of Biological Chemistry. 2004;279: 45546– 45555. 10.1074/jbc.M402762200

42. Keyamura K, Fujikawa N, Ishida T, Okazaki S, Su’etsugu M, Fujimitsu K, et al. The interaction of DiaA and DnaA regulates the replication cycle in E. coli by directly promoting ATP DnaA-specific initiation complexes. Genes Dev. 2007; 2083–2099. Available: 10.1101/gad.1561207

43. Sugiyama Y, Nakamura A, Matsumoto M, Kanbe A, Sakanaka M, Higashi K, et al. A Novel Putrescine Exporter SapBCDF of Escherichia coli. Journal of Biological Chemistry. 2016;291: 26343–26351. 10.1074/jbc.M116.762450

44. Al Mamun AAM, Lombardo M-J, Shee C, Lisewski AM, Gonzalez C, Lin D, et al. Identity and Function of a Large Gene Network Underlying Mutagenic Repair of DNA Breaks. Science (1979). 2012;338: 1344–1348. doi:10.1126/science.1226683

45. T BE, Debbie M, Josef N, Wolfgang E. Multiple Paths for Nonphysiological Transport of K+ in Escherichia coli. J Bacteriol. 2004;186: 4238–4245. doi:10.1128/jb.186.13.4238-4245.2004

46. Tscherne JS, Nurse K, Popienick P, Michel H, Sochacki M, Ofengand J. Purification, Cloning, and Characterization of the 16S RNA m5C967 Methyltransferase from Escherichia coli. Biochemistry. 1999;38: 1884–1892. doi:10.1021/bi981880l

47. Toussaint A, Faelen M. The dependence of temperate phage mu-1 upon replication functions of E. coli K12. Mol Gen Genet. 1974;131: 209–214. doi:10.1007/BF00267960

48. Erzberger JP, Mott ML, Berger JM. Structural basis for ATP-dependent DnaA assembly and replication-origin remodeling. Nat Struct Mol Biol. 2006;13: 676– 683. doi:10.1038/nsmb1115

49. Speck C, Weigel C, Messer W. ATP– and ADP–DnaA protein, a molecular switch in gene regulation. EMBO J. 1999;18: 6169–6176–6176. 10.1093/emboj/18.21.6169

50. Yung BY, Kornberg A. Membrane attachment activates dnaA protein, the initiation protein of chromosome replication in Escherichia coli. Proceedings of the National Academy of Sciences. 1988;85: 7202–7205. doi:10.1073/pnas.85.19.7202

51. Crooke E, Castuma CE, Kornberg A. The chromosome origin of Escherichia coli stabilizes DnaA protein during rejuvenation by phospholipids. Journal of Biological Chemistry. 1992;267: 16779–16782. 10.1016/S0021-9258(18)41849-2

52. Zheng W, Li Z, Skarstad K, Crooke E. Mutations in DnaA protein suppress the growth arrest of acidic phospholipid&#x2010;deficient *Escherichia coli* cells. EMBO J. 2001;20: 1164–1172–1172. 10.1093/emboj/20.5.1164

53. Li Z, Kitchen JL, Boeneman K, Anand P, Crooke E. Restoration of Growth to Acidic Phospholipid-deficient Cells by DnaA(L366K) Is Independent of Its Capacity for Nucleotide Binding and Exchange and Requires DnaA. Journal of Biological Chemistry. 2005;280: 9796–9801. doi:10.1074/jbc.M413923200

54. Hou Y, Kumar P, Aggarwal M, Sarkari F, Wolcott KM, Chattoraj DK, et al. The linker domain of the initiator DnaA contributes to its ATP binding and membrane association in E. coli chromosomal replication. Sci Adv. 2024;8: eabq6657. doi:10.1126/sciadv.abq6657

55. Su’etsugu M, Shimuta T, Ishida T, Kawakami H, Katayama T. Protein Associations in DnaA-ATP Hydrolysis Mediated by the Hda-Replicase Clamp Complex. Journal of Biological Chemistry. 2005;280: 6528–6536. 10.1074/jbc.M412060200

56. Dalrymple BP, Kongsuwan K, Wijffels G, Dixon NE, Jennings PA. A universal protein–protein interaction motif in the eubacterial DNA replication and repair systems. Proceedings of the National Academy of Sciences. 2001;98: 11627– 11632. doi:10.1073/pnas.191384398

57. Mathee K, Howe MM. Identification of a positive regulator of the Mu middle operon. J Bacteriol. 1990;172: 6641–6650. doi:10.1128/jb.172.12.6641-6650.1990

58. Mathee K, Howe MM. Bacteriophage Mu Mor protein requires sigma 70 to activate the Mu middle promoter. J Bacteriol. 1993;175: 5314–5323. doi:10.1128/jb.175.17.5314-5323.1993

59. Kumaraswami M, Howe MM, Park H-W. Crystal Structure of the Mor Protein of Bacteriophage Mu, a Member of the Mor/C Family of Transcription Activators. Journal of Biological Chemistry. 2004;279: 16581–16590. 10.1074/jbc.M313555200

60. Jang S, Harshey RM. Repair of transposable phage Mu DNA insertions begins only when the . coli replisome collides with the transpososome. Mol Microbiol. 2015;97: 746–758. 10.1111/mmi.13061

61. Jang S, Sandler SJ, Harshey RM. Mu Insertions Are Repaired by the Double-Strand Break Repair Pathway of Escherichia coli. PLoS Genet. 2012;8: e1002642–2 Available: 10.1371/journal.pgen.1002642

62. Guo L, Katayama T, Seyama Y, Sekimizu K, Miki T. Isolation and characterization of novel cold-sensitive dnaA mutants of Escherichia coli. FEMS Microbiol Lett. 1999;176: 357–366. doi:10.1111/j.1574-6968.1999.tb13684.x

63. Takata M, Guo L, Katayama T, Hase M, Seyama Y, Miki T, et al. Mutant DnaA proteins defective in duplex opening of oriC, the origin of chromosomal DNA replication in Escherichia coli. Mol Microbiol. 2000;35: 454–462. 10.1046/j.1365-2958.2000.01722.x

64. Garner J, Durrer P, Kitchen J, Brunner J, Crooke E. Membrane-mediated Release of Nucleotide from an Initiator of Chromosomal Replication, Escherichia coli DnaA, Occurs with Insertion of a Distinct Region of the Protein into the Lipid Bilayer. Journal of Biological Chemistry. 1998;273: 5167–5173. 10.1074/jbc.273.9.5167

65. Makise M, Mima S, Tsuchiya T, Mizushima T. Identification of Amino Acids Involved in the Functional Interaction between DnaA Protein and Acidic Phospholipids. Journal of Biological Chemistry. 2000;275: 4513–4518. 10.1074/jbc.275.6.4513

66. Parks AR, Li Z, Shi Q, Owens RM, Jin MM, Peters JE. Transposition into Replicating DNA Occurs through Interaction with the Processivity Factor. Cell. 2009;138: 685–695. 10.1016/j.cell.2009.06.011

67. Kritaya K, Peter J, J PM, Gene W, Brian D. The Plasmid RK2 Replication Initiator Protein (TrfA) Binds to the Sliding Clamp β Subunit of DNA Polymerase III: Implication for the Toxicity of a Peptide Derived from the Amino-Terminal Portion of 33-Kilodalton TrfA. J Bacteriol. 2006;188: 5501–5509. doi:10.1128/jb.00231-06

68. D KP, Trevor B, M LD, C TJ, Eltz CJ, Elliot C, et al. Identification of a Novel Membrane-Associated Gene Product That Suppresses Toxicity of a TrfA Peptide from Plasmid RK2 and Its Relationship to the DnaA Host Initiation Protein. J Bacteriol. 2003;185: 1817–1824. doi:10.1128/jb.185.6.1817-1824.2003

69. Millar D, Trakselis MA, Benkovic SJ. On the Solution Structure of the T4 Sliding Clamp (gp45). Biochemistry. 2004;43: 12723–12727. doi:10.1021/bi048349c

70. Moarefi I, Jeruzalmi D, Turner J, O’Donnell M, Kuriyan J. Crystal structure of the DNA polymerase processivity factor of T4 bacteriophage11Edited by I. A. Wilson. J Mol Biol. 2000;296: 1215–1223. 10.1006/jmbi.1999.3511

71. Gibson DG, Young L, Chuang R-Y, Venter JC, Hutchison CA, Smith HO. Enzymatic assembly of DNA molecules up to several hundred kilobases. Nat Methods. 2009;6: 343–345. doi:10.1038/nmeth.1318

72. Datsenko KA, Wanner BL. One-step inactivation of chromosomal genes in *Escherichia coli* K-12 using PCR products. Proceedings of the National Academy of Sciences. 2000;97: 6640–6645. doi:10.1073/pnas.120163297

73. Ronda C, Pedersen LE, Sommer MOA, Nielsen AT. CRMAGE: CRISPR Optimized MAGE Recombineering. Sci Rep. 2016;6: 19452. doi:10.1038/srep19452

74. A KJ, G SA, T LM. The Stringent Response Inhibits DNA Replication Initiation in E. coli by Modulating Supercoiling of oriC. mBio. 2019;10: 10.1128/mbio.01330-19. doi:10.1128/mbio.01330-19

75. Altschul SF, Madden TL, Schäffer AA, Zhang J, Zhang Z, Miller W, et al. Gapped BLAST and PSI-BLAST: a new generation of protein database search programs. Nucleic Acids Res. 1997;25: 3389–3402. doi:10.1093/nar/25.17.3389

